# Transformation of neural coding for vibrotactile stimuli along the ascending somatosensory pathway

**DOI:** 10.1101/2023.10.13.562210

**Authors:** Kuo-Sheng Lee, Alastair Loutit, Dominica de Thomas Wagner, Mark Sanders, Mario Prsa, Daniel Huber

## Abstract

Perceiving substrate vibrations is a fundamental component of somatosensation. In mammals, action potentials fired by rapidly adapting mechanosensitive afferents are known to reliably time lock to the cycles of a vibration. This stands in contrast to coding in the higher-order somatosensory cortices, where neurons generally encode vibrations in their firing rates, which are tuned to a preferred vibration frequency. How and where along the ascending neuraxis is the peripheral afferent temporal code of cyclically entrained action potentials transformed into a rate code is currently not clear. To answer this question, we probed the encoding of vibrotactile stimuli with electrophysiological recordings along major stages of the ascending somatosensory pathway in mice. Recordings from individual primary sensory neurons in lightly anesthetized mice revealed that rapidly adapting mechanosensitive afferents innervating Pacinian corpuscles display phase-locked spiking for vibrations up to 2000 Hz. This precise temporal code was reliably preserved through the brainstem dorsal column nuclei. The main transformation step was identified at the level of the thalamus, where we observed a significant loss of phase-locked spike timing information accompanied by a further narrowing of tuning curve widths. Using optogenetic manipulation of thalamic inhibitory circuits, we found that parvalbumin-positive interneurons in thalamic reticular nucleus participate in sharpening frequency selectivity and disrupting the precise spike timing of ascending neural signals encoding vibrotactile stimuli. To test the functional implications of these different neural coding mechanisms, we applied frequency-specific microstimulation within the brainstem, which generated frequency selectivity reminiscent of real vibration responses in the somatosensory cortex, whereas microstimulation within thalamus did not. Finally, we applied microstimulation in the brainstem of behaving mice and demonstrated that frequency-specific stimulation could provide informative and robust signals for learning. Taken together, these findings not only reveal novel features of the computational circuits underlying vibrotactile sensation, but could also guide biomimetic stimulus strategies to activate specific nuclei along the ascending somatosensory pathway for sensory neural prostheses.

## Introduction

Vibrations are a fundamental component of our tactile experience. We use our ability to sense vibrations in many daily situations, such as perceiving incoming phone calls or detecting an approaching vehicle. In mammals, high frequency vibrations (>100Hz) are transduced to electrical signals by Pacinian corpuscles (PC), which are innervated by primary sensory neurons whose soma are situated in the dorsal root ganglia (DRG). This information is relayed to second order neurons in the dorsal column nuclei (DCN, gracile and cuneate) of the medulla. DCN axons project to neurons of the ventral posterolateral nucleus of the thalamus (VPL), which in turn project to primary somatosensory cortex (S1). The earliest electrophysiological recordings from receptor end organs (Adrian and Umrath 1929; Mountcastle, LaMotte, and Carli 1972) revealed that the receptor potentials, as well as the resulting action potentials in DRG neurons, reliably time-lock to the cycles of a vibration up to 1000 Hz. However, little is known about how this precise temporal coding is processed in the rest of the central nervous system to achieve feature specific computation, commonly with a rate code.

In the medulla, only very few studies in cats have recorded neuronal responses to vibration, and they reported mostly 1:1 locking of spikes to the phase of the stimulus in low frequencies (Gynther, Vickery, and Rowe 1995; Rowe 2002; Sahai et al. 2006). In contrast, VPL has been more extensively studied and is considered an integration hub for ascending somatosensory information along the lemniscal pathway. Individual neurons were found to be occasionally temporally entrained, and there was an increasing complexity in preferred frequencies and neural adaptation to vibration in this area (Herron and Dykes 1986; Vázquez, Salinas, and Romo 2013).

Finally, at the cortical level different coding modes have been described. On one hand, in primate S1, vibration information seems to be represented by precise spiking patterns (Suresh, Saal, and Bensmaia 2016; Saal, Harvey, and Bensmaia 2015; Suresh et al. 2021), similar to the cutaneous mechanoreceptive afferents (Birznieks et al. 2019; Birznieks and Vickery 2017). It is not until the secondary somatosensory cortex (S2), that the frequency-dependent firing rate modulations become the primary neural code for vibration and phase-locked responses seem absent (Salinas et al. 2000). On the other hand, in rodent S1, neurons seem to represent vibration frequency by a rate code tuned to a preferred stimulus feature (Prsa et al. 2019). This resembles feature-selective coding found in many other sensory modalities (Prsa et al. 2019). Yet it raises an important question: how are cyclically entrained action potentials (a temporal code) in afferents of peripheral mechanoreceptors transformed along the ascending neuraxis into a rate code in the cortex? Although a similar transformation of low frequencies (< 10 Hz) has been investigated in the rat whisker system (Ahissar, Sosnik, and Haidarliu 2000), it remains unclear for high frequency encoding of skin vibrotactile stimuli. In the current study, we addressed this question in mice by recording along the dorsal column medial lemniscus pathway to first uncover the basic coding principles and rules underlying sensory perception. Then, we sought to uncover the circuit mechanism underlying the transformation of neural coding with optogenetic manipulations. Finally, we tested to what extent these coding strategies might be leveraged as artificial sensory feedback in neuroprosthetics using microstimulation to selected neuronal populations. Overall, our results provide key insights into how to artificially mimic vibrotactile stimuli in neuroprosthetic devices.

## Results

### Temporal code of vibrotactile stimuli along the ascending pathway

To identify the location and mechanism of the transformation of neural coding for vibration, we first conducted in vivo extracellular recordings in anesthetized mice along the ascending somatosensory pathway (**Figure 1a**). Pure sinusoidal vibrations of a wide range of frequencies (100 Hz to 1900 Hz) (Prsa et al. 2019) were delivered to the hindlimb of anesthetized mice. Frequency and amplitude combinations were selected based on previous psychophysical and physiological findings, in which mice can perceive stimuli as gentle vibration (K.-S. Lee et al. 2023; Prsa et al. 2019, 2021). We computed standardized inter-spike intervals (ISIs; see **Methods**) whose distribution should peak at integer multiples of the stimulus period in the case of phase-locked entrainment of action potentials by the oscillatory stimulus (**Figure 1b-e and Figure S1**). Nerve recordings from low-threshold mechanoreceptors that are likely innervating Pacinian corpuscles (Aβ RA2-LTMR; **Figure S2**), display phase-locked spiking for vibrations up to 1900 Hz (**Figure 1b**). We show that the precise temporal code found in the periphery is surprisingly well preserved in putative second-order neurons in the brainstem gracile nucleus (**Figure 1c**). This reliable high-frequency spiking transmission in DCN likely relies on specialized mechanisms, possibly endocytic processes and synaptic structures (Wu, Ryan, and Lagnado 2007). We therefore conducted reconstruction of focused ion beam scanning electron microscopy data of DCN synapses and expected to find structures similar to either the ribbon synapses in cochlear hair cells (Lenzi et al. 1999), or the calyx of Held in the auditory brainstem (Sun, Wu, and Wu 2002). However, we did not find evidence of either, suggesting that different mechanisms might govern high-throughput transmission in DCN (**Figure S3**), consistent with previous observations with transmission electron microscopy (Ellis and Rustioni 1981; Banna and Jabbur 1989).

**Figure 1.**
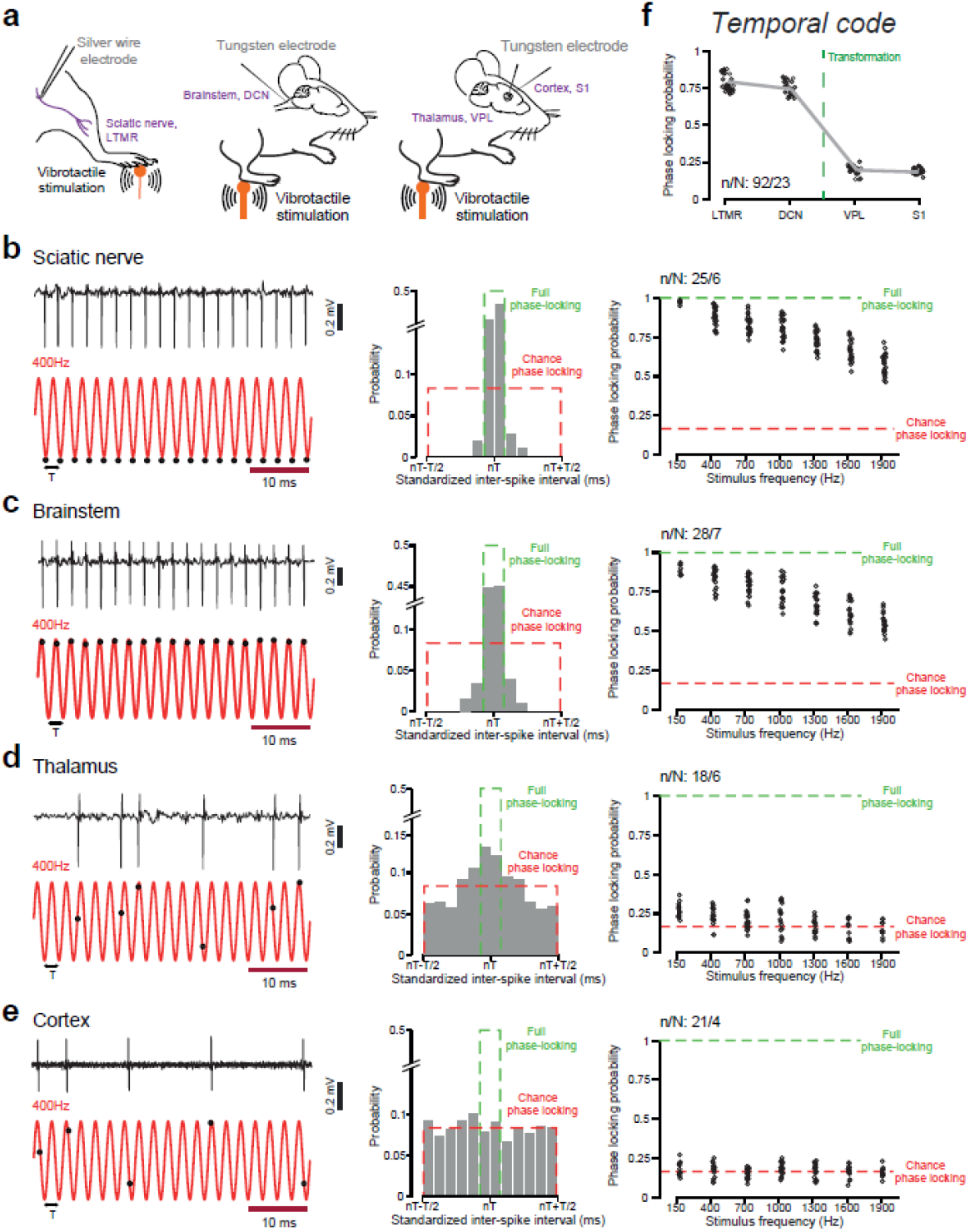
Loss of temporal coding for vibration beyond the brainstem in the ascending pathway. **a**, *In vivo* experimental setup schematic in anesthetized animals. Left, nerve fiber recording of sciatic nerve. Middle, recording of brainstem from the back of the neck. Right, recording of thalamus and cortex. **b**, Left, representative trace of a mechanosensitive afferent spiking in response to 400 Hz vibration stimulus. The timing of its action potentials (black dots) relative to periodic cycles of the vibration (red) shows nearly perfect entrainment. Middle, distribution of standardized inter-spike intervals of an example neuron at this frequency. Right, phase-locking probability measured for n = 25 neurons (6 mice) at each tested frequency (black circles). **c-e**, Same as in **b**, for brainstem, thalamus and cortex, respectively. f, The phase-locking probability compared between all four areas in the ascending pathway by averaging the results tested from 100 to 1900 Hz (Kruskal–Wallis test, P < 0.0001). Dunn’s Test: (S1=VPL)<(LTMR=DCN).

In contrast to the reliable transmission revealed at the DCN level, precise temporal coding is lost in thalamic VPL (**Figure 1d**) and remains absent in somatosensory cortex (**Figure 1e**). Phase-locking probability thus suddenly drops downstream of the brainstem (**Figure 1f**), suggesting that loss of temporal coding for vibration in the ascending pathway occurs at the level of the thalamus.

### Rate code of vibrotactile stimuli in the ascending pathway

The absence of cyclically phase-locked spiking beyond the brainstem suggests that vibration frequencies are represented as pure rate codes in both thalamus and cortex (Prsa et al. 2019). However, rate coding for different vibration frequencies could also exist at early stages of the ascending pathway. We therefore investigated the feature-selective encoding of substrate vibrations within the same data as in **Figure 1**. Most of the vibration responsive units, even from nerve fibers, showed a frequency preference across various amplitudes (**Figure 2a and Figure S4**). Nerve recordings revealed broad tuning curves covering a wide range of preferred frequencies (**Figure 2b**), which indicated a moderate level of rate coding for vibration in the periphery. The pattern and distribution of tuning curves in the brainstem gracile nucleus were similar to the periphery (**Figure 2c**). In contrast, tuning curves in the ventral posterolateral nucleus of the thalamus became much more narrowly tuned (**Figure 2d**), as did those in somatosensory cortex (**Figure 2e**). Comparison of tuning curve widths for vibration frequency, defined by the half-width at half-maximum amplitude (HWHM), from the periphery to cortex (**Figure 2f**), suggests that the main step causing a narrowing of tuning found in S1 involves thalamic circuits. In addition, we found the coefficient of variation (CV) of inter-spike intervals also greatly increased at the thalamus (mean CV: LTMR, 0.019; DCN, 0.024; VPL, 0.975; S1, 1.088). Taken together, our results show that the neural code transformation, from mechanoreceptor temporal encoding to cortical rate encoding, thus mainly occurs in thalamus (**Figure 2g**).

**Figure 2.**
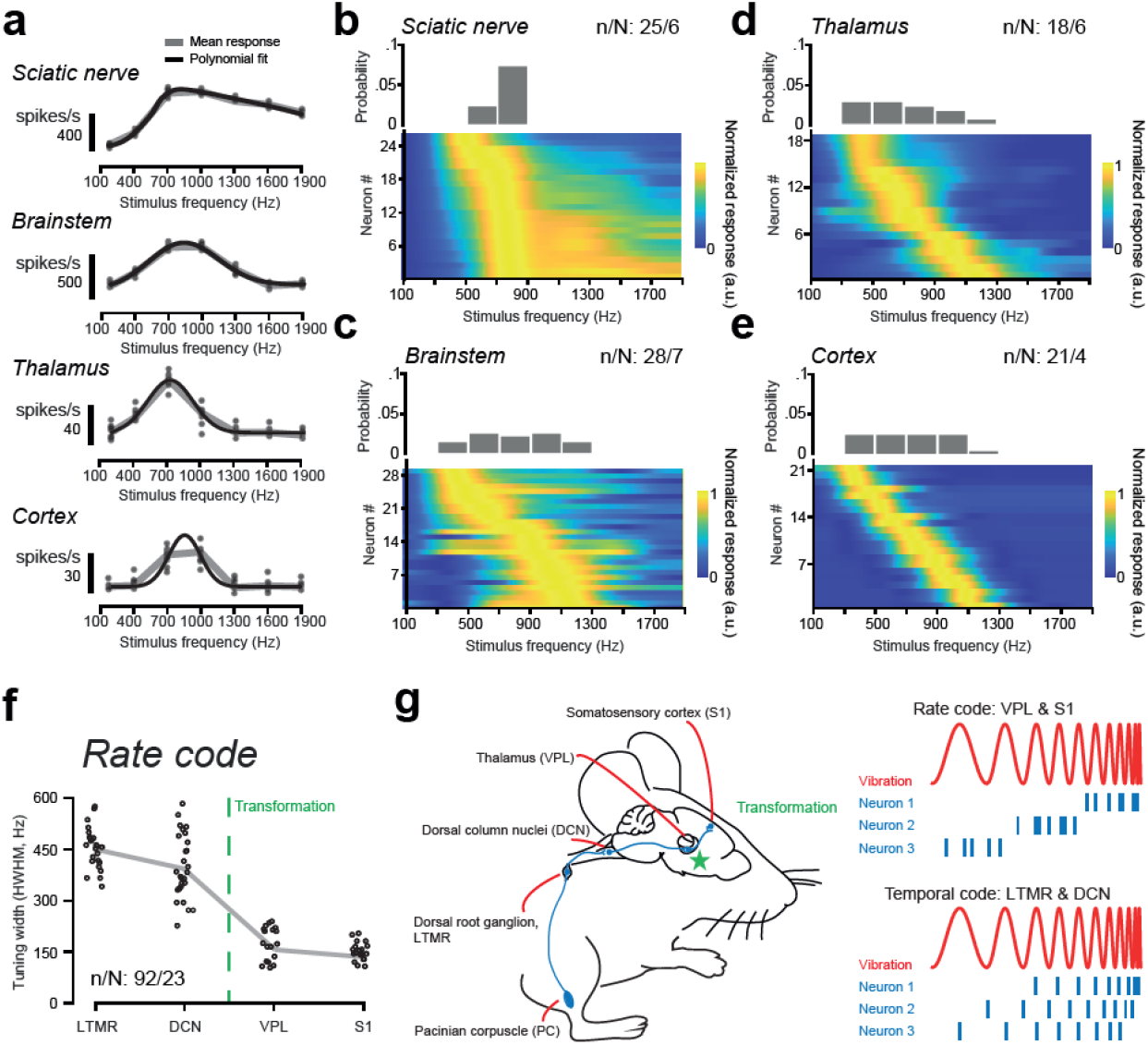
Evolution of rate code for vibration of the ascending pathway. **a**, Tuning curves based on mean spike rates for example units recorded from peripheral nerve, brainstem, thalamus and cortex, respectively. **b**, Normalized tuning curves of all units from sciatic nerve recordings with identifiable tuning curve peaks sorted by decreasing best frequency and distribution of neurons’ best frequencies tested at 5 µm amplitude. **c-e**, Similar as in **b**, for brainstem, thalamus and cortex, respectively. **f**, The tuning curve, half-width at the half-maximum of the curve (HWHM), for vibration frequency compared between all four areas in the ascending pathway (Kruskal–Wallis test, P < 0.0001). Dunn’s Test: (S1=VPL)>(LTMR=DCN). **g**, A model for transformation of neural codes in the hindlimb dorsal column–medial lemniscal pathway.

### The role of inhibitory thalamic circuits

According to previous work, loss of temporal coding can be partially attributed to strong, frequency-dependent depressing-type excitatory connections, which supply ‘driver’ type inputs (Sherman and Guillery 1998) between DCN and VPL (Yamawaki et al. 2021). However, this would not explain the significant narrowing of rate code tuning (**Figure 2**). This aspect might be better explained by local interconnected feedforward and feedback circuits (Temereanca and Simons 2004; Briggs and Usrey 2011; Pinault and Deschênes 1998; Y.-T. Li et al. 2013). To disentangle how the neural circuits within the thalamus could contribute to the transformation of the neural coding, we manipulated the neuronal activity with optogenetic tools. It has been shown that parvalbumin positive inhibitory interneurons participate in improving feature selectivity in visual cortex (S.-H. Lee et al. 2012) and controlling temporal firing patterns in spinal dorsal horn (Rankin et al. 2023). In rodents, sensory thalamus receives inhibitory drive from the thalamic reticular nucleus (Martinez-Garcia et al. 2020; Y. Li et al. 2020). Here, we paired in vivo optogenetic stimulation through fiber optics with extracellular recordings in thalamic nuclei (**Figure 3 and Figure S5**).

**Figure 3.**
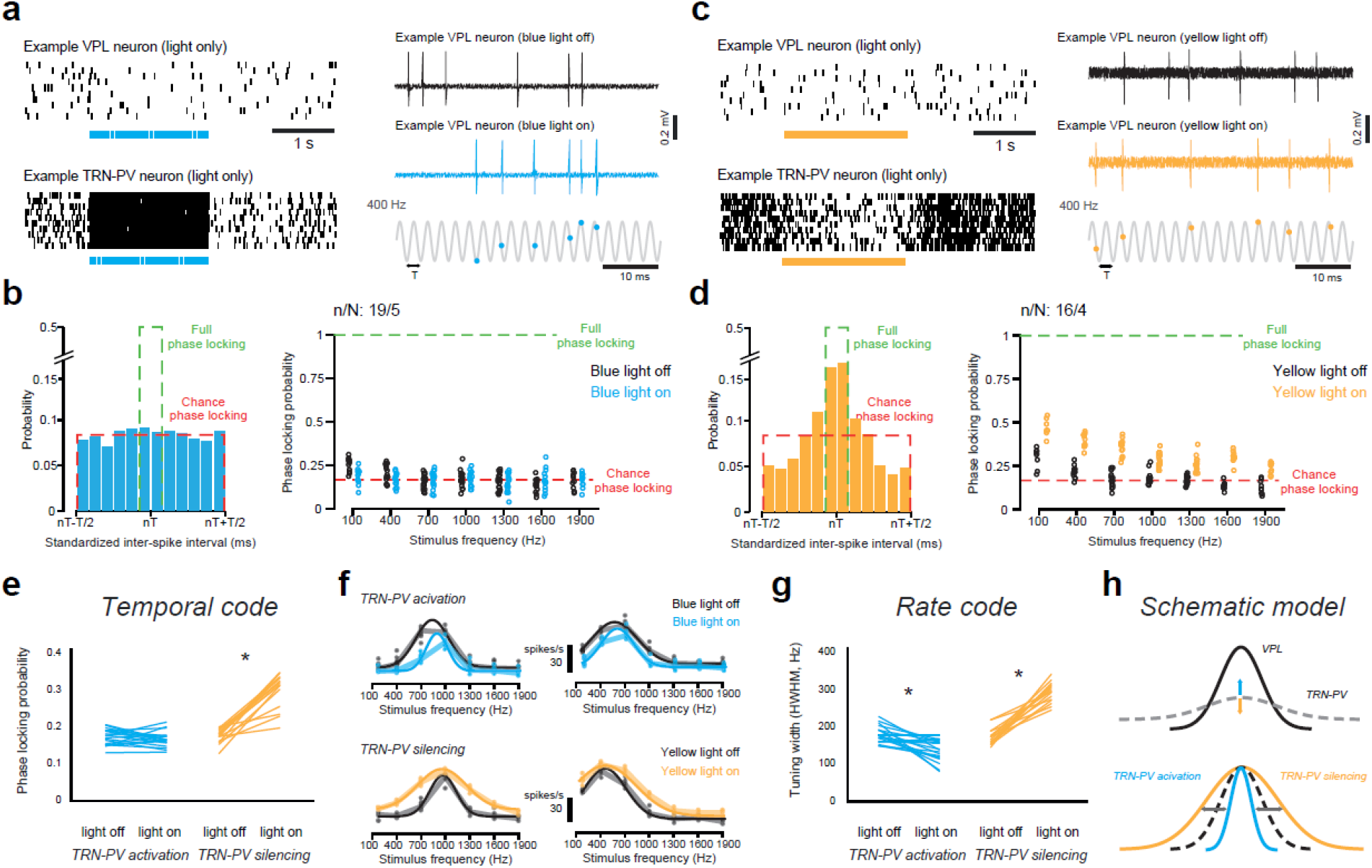
Bi-directional optogenetic manipulation of parvalbumin neurons in thalamic reticular nucleus (TRN-PV) reveals its contribution to neural coding. **a**, Example recording of a VPL neuron (left top) and a TRN-PV neuron (left bottom) in response to 40 Hz (1ms pulses) of 473 nm blue light stimulation at TRN, in *PV-Cre* × *ChR2*-EYFP (Ai32) mice. Right, example recording of a VPL neuron in response to 400 Hz vibration at the hindlimb, with light off (black) or with light on (blue). The timing of its action potentials (blue dots) relative to periodic cycles of the 400-Hz vibration (gray) shows no sign of entrainment. **b**, Left, with blue light stimulation, the distribution of standardized inter-spike intervals of an example VPL neuron at 400 Hz vibration is nearly at random entrainment. Right, phase-locking probability measured for n = 16 VPL neurons (6 mice) at each tested frequency evoking significant responses, either with light off (black circles) or with light on (blue circles). **c**, Example recording of a VPL neuron (left top) and a TRN-PV neuron (left bottom) in response to 2s continuous 568 nm yellow light stimulation at TRN, in *PV-Cre* × *ArchT*-GFP (Ai40) mice. Right, example recording of a VPL neuron in response to 400 Hz vibration at the hindlimb, with light off (black) or with light on (yellow). The timing of its action potentials (yellow dots) relative to periodic cycles of the 400-Hz vibration (gray) shows little entrainment. **d**, Left, with yellow light stimulation, the distribution of standardized inter-spike intervals of an example VPL neuron at 400 Hz vibration is slightly above chance entrainment. Right, phase-locking probability measured for n = 17 VPL neurons (6 mice) at each tested frequency, either with light off (black circles) or with light on (yellow circles). **e**, The phase-locking probability of VPL neurons affected by optogenetic activation or inactivation of TRN-PV neurons (paired-sample t-test, *P < 0.0001). **f**, Top, tuning curve sharpening of 2 VPL example neurons after TRN-PV activation by blue light. Bottom, tuning curve broadening of 2 VPL example neurons after TRN-PV inactivation by yellow light. g, Academia Sinica. h, Schematic model showing how TRN-PV neurons sharpen the tuning curve of VPL neurons.

To achieve bi-directional optogenetic manipulation of parvalbumin neurons in the thalamic reticular nucleus (TRN-PV), we selectively expressed both excitatory Channelrhodopsin-2 (ChR2) and inhibitory Archaerhodopsin-3 (ArchT) in TRN-PV by using a double-transgenic strategy (PV-Cre × ChR2-EYFP, PV-Cre × ArchT-GFP, **Figure S5**). We first recorded TRN-PV and VPL neuron responses to blue light stimulation and to vibration stimuli paired with this optogenetic activation (**Figure 3a**). We found that optogenetic activation of TRN-PV neurons did not alter temporal processing of vibration in VPL (**Figure 3b**). However, inactivation of TRN-PV neurons in PV-Cre × ArchT-GFP mice, via yellow light stimulation (**Figure 3c**),contributed to the phase-locking probability of VPL neurons (**Figure 3d**). Note that the phase-locking probability never reached that of the brainstem level, which suggests that other mechanisms such as intrinsic synaptic properties (Sherman and Guillery 1998)(S.-H. Lee et al. 2012; Y.-T. Li et al. 2012) (Yamawaki et al. 2021)(Sherman and Guillery 1998) must also contribute to the loss of the temporal code. Furthermore, we demonstrated that TRN-PV neurons are generally broadly tuned to vibration, and their inactivation can broaden VPL neuron tuning widths (**Figure 3g, and Figure S5**). We deduced that TRN-PV neuronal activity sharpens VPL neuron frequency selectivity and contributes to the loss of temporal coding (**Figure 3e, f**). Thus, the mechanism by which they achieve this is similar to the broad inhibition that sharpens feature selectivity in the visual cortex (S.-H. Lee et al. 2012; Y.-T. Li et al. 2012) (**Figure 3h**).

### Probing the neural coding mechanisms with microstimulation

To explicitly test if the coding mechanism described above could be used to artificially drive neurons in the somatosensory system (Flesher et al. 2016), we probed frequency selective neuron responses in S1 during temporally structured microstimulation applied to the DCN and VPL (**Figure 4a and Figure S6**) (Stieger et al. 2022). First, the vibration frequency tuning of S1 neurons was determined using two photon calcium imaging in anesthetized CaMKII-tTA × TRE-GCaMP6s mice (see **Methods**). Subsequently, we characterized responses of the same S1 neurons to electrical stimulation (bipolar platinum–iridium electrodes) using matched frequencies of electrical pulses applied to the DCN (**Figure 4b**). Microstimulation resulted in frequency selective tuning curves comparable to the tuning curves evoked by hindlimb vibration (**Figure 4c**). To compare the responses of both modalities we calculated two indices: a “temporal coding index” (TCI, correlation of frequency tuning curves for vibration and electrical stimulation) and “rate coding index” (RCI, correlation of tuning curves to frequency invariant responses depending only on the number of pulses, see **Methods**). Equivalent experiments were then conducted with VPL stimulation (**Figure 4d-e**). TCI and RCI show that the neural coding within DCN and VPL can be fully distinguished as a non-overlapping distribution (**Figure 4f**), which could be partly attributed to the heterogeneity of VPL response properties, and the amount of convergence/divergence in thalamic circuits (Ahissar, Sosnik, and Haidarliu 2000). To further support this idea, microstimulation within the spinal dorsal column paired with frequency selectivity readout at DCN also showed that feedforward innervation from the periphery is mostly based on a temporal code (**Figure S7**). In conclusion, temporally precise microstimulation within the DCN, but not within the VPL, can reproduce S1 frequency tuning for vibration, confirming significant differences in the neural coding mechanisms in these two areas (**Figure 4g**).

**Figure 4.**
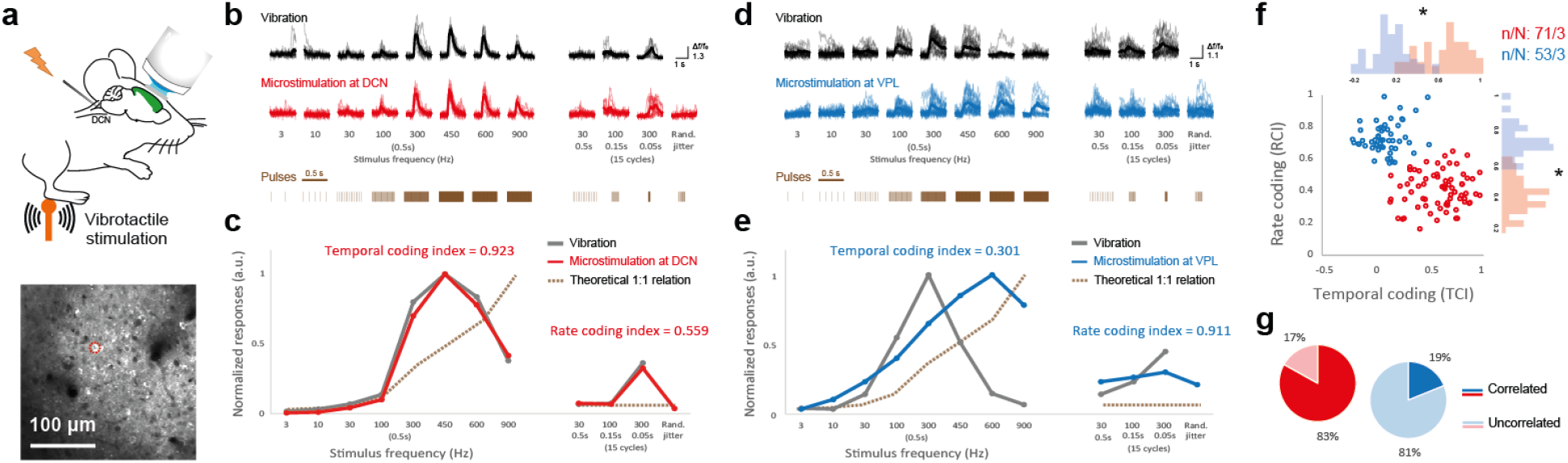
Microstimulation reveals different neural codes for vibration in the brainstem and thalamus. **a,** Experimental setup for two-photon calcium imaging in somatosensory cortex with *CaMKII-tTA* × *TRE*-GCaMP6s mice (top) and an example field-of-view (bottom). **b,** Top, averaged relative changes in fluorescence (Δf/f0) of an example S1 neuron imaged with two-photon microscopy in response to vibrations of different frequencies and durations. Bottom, averaged Δf/f0 of the same neuron imaged with two-photon microscopy in response to contralateral DCN microstimulation of different frequencies and durations. Background lines are responses to individual trials. The structure of the pulses was displayed in a different scale for visualization. **c,** Vibration frequency-selective tuning curve (gray) and DCN microstimulation frequency-selective tuning curves (red) of the example neuron in b. Gray dashed line, hypothetical response pattern predicted from the number of electrical pulses, assuming that the neuron increases its firing rate linearly. Temporal coding index (TCI) and rate coding index (RCI) were computed from the tuning curves (see Methods). **d,** Top, averaged Δf/f0 of an example S1 neuron imaged with two-photon microscopy in response to vibrations of different frequencies and durations. Bottom, averaged Δf/f0 of the same neuron imaged with two-photon microscopy in response to ipsilateral VPL microstimulation of different frequencies and durations. Background lines are responses to individual trials. **e,** Vibration frequency-selective tuning curve (gray) and VPL microstimulation frequency-selective tuning curves (blue) of the example neuron in d. gray dashed line, hypothetical response pattern predicted from the number of electrical pulses. **f,** The distribution of temporal coding index (TCI) and rate coding index (RCI) for both DCN stimulation (red) and VPL stimulation (blue) showed very different coding strategies (unpaired t-test, *P < 0.0001). **g,** The percentage of the S1 neurons whose vibration tuning curve can be reproduced by microstimulation tuning curves (Pearson’s correlation).

### Perceptual relevance of temporal coding in the brainstem

Finally, we tested if DCN stimulation with high temporal precision could be perceived and thus potentially used for biomimetic feedback in neuroprosthetic devices. Because of the dense passage of fibers and the close proximity of nuclei involved in vital functions, such interventions have not yet been attempted. We therefore developed a new chronic stimulation method that allows us to target the DCN unilaterally (see **Methods**). Inspired by other cortical microstimulation tasks (Romo et al. 1998; Callier et al. 2020), we trained mice (n = 5) to perform a frequency discrimination task from artificial sensory stimulation interleaved between vibration stimuli applied to the hind limbs (**Figure 5a**, see **Methods**). All five mice learned to discriminate 200 Hz from 800 Hz vibration stimuli applied to the hindlimbs (**Figure 5b**) and were then, on test days, challenged to discriminate 200 Hz from 600 Hz electrical microstimulation applied to either the DCN or S1 on 20% of the trials, while the other 80% of trials remained vibrations applied to the hindlimb (**Figure 5a,c)**. On the first day that electrical stimulation was introduced, performance for electrical stimulation trials was at chance level for 4/5 animals. Two animals experienced DCN microstimulation first and three animals experienced S1 microstimulation first. After training with electrical microstimulation in both regions (**Figure 5d**), all five animals performed significantly better for DCN microstimulation compared to S1, suggesting mice were better able to relate the DCN electrical stimulation frequencies to the vibrations of the same frequency. These results are reinforced by the discrimination of only electrical stimuli by the same five mice as performance in response to DCN stimulation was significantly better than for S1 (**Figure 5e** and legend). Most of the electrodes continued functioning even after two months of implantation (**Figure S8**). These results establish for the first time that mice can be chronically implanted with DCN electrodes by which frequency-specific stimulation can provide informative and robust signals for learning, which in most cases were better than cortical microstimulation in S1 (**Figure 5c**).

**Figure 5.**
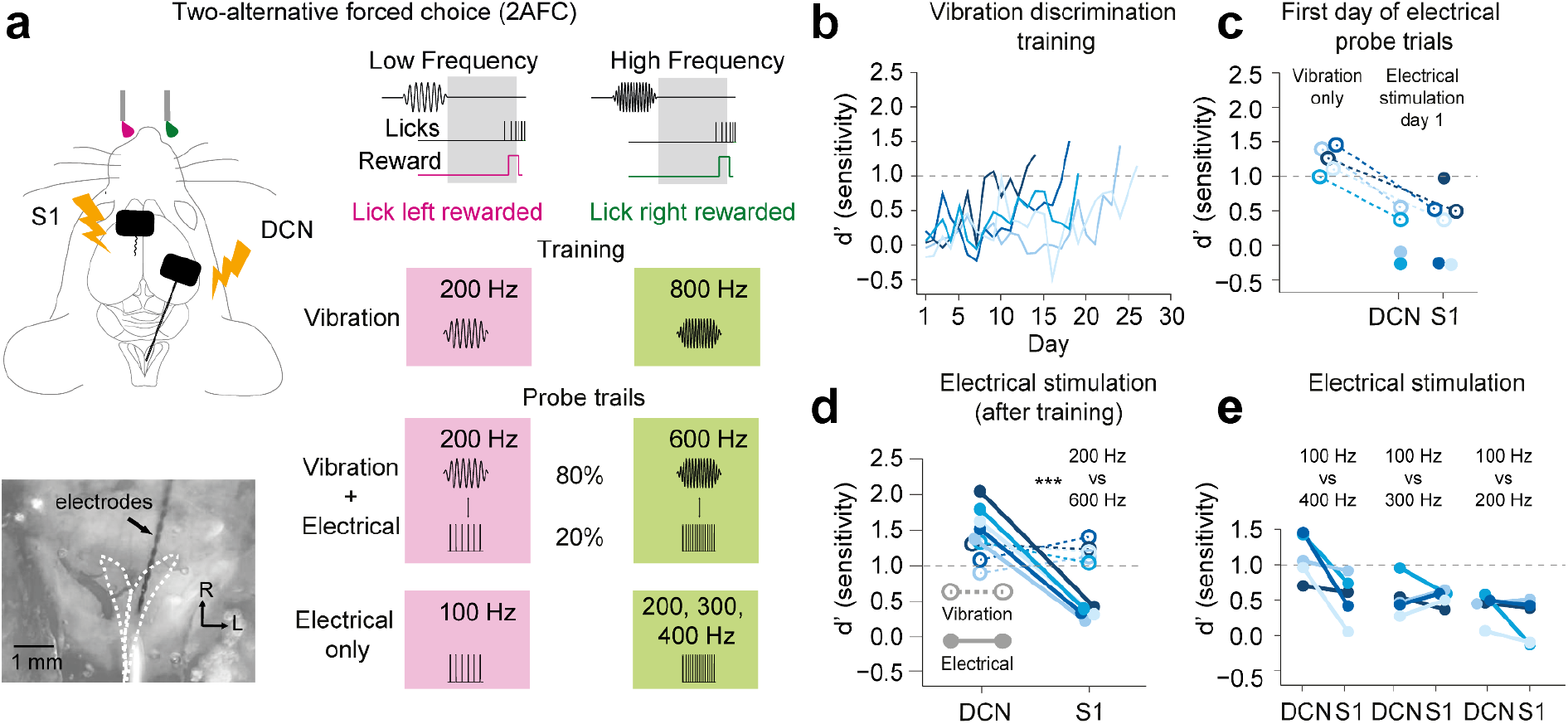
Discrimination of vibration and artificial sensory microstimulation. **a,** Schematic showing the two-alternative forced choice task (top left), with a dorsal view of the dorsal column nuclei (DCN) during electrode implantation (bottom left). White dashed lines indicate the boundaries of the left and right gracile nuclei. Mice were trained to discriminate vibration stimulation frequency pairs applied to the hindpaws. Comparison frequencies were 200 Hz and 800 Hz. Mice learned to discriminate 200 Hz and 800 Hz during training, and this was reduced to 200 Hz compared to 600 Hz during test days for the Vibration + Electrical stimulation task. Electrical stimulus probe trials of the same frequencies as for vibrations were introduced in 20% of stimulus presentations, randomly interleaved with the vibration trials. Electrical stimuli were applied to either the right DCN or left somatosensory cortex hindlimb region (S1). For probe trials, the licking choice task was the same as for training on vibration only. **b,** Learning curves represented as d-prime (d’) calculated from hits and misses from mice discriminating between 200 Hz and 800 Hz hindlimb vibration stimulation, over training days. Testing commenced after mice reached d’ > 1 (gray dashed line) over 100 consecutive trials. **c,** D’ values showing how performance changed from the end of training on vibration stimuli (200 Hz vs 800 Hz) to the first day that mice experienced the interleaved electrical microstimulation trials (200 Hz vs 600 Hz). Two mice experienced DCN electrical microstimulation first, while the other three experienced S1 stimulation first. **d,** Performance on the same task as in c after mice had been trained on electrical stimulation in both DCN and S1. After all mice had experienced electrical stimulation to both DCN and S1, d’ values were significantly greater for electrical stimulation applied to DCN compared to S1 (\*\*\**p* = 0.0001, *t* = 15.4, paired *t*-test). **e,** In another task, the same mice discriminated only electrical stimuli in which they were rewarded for licking right for either 200, 300, and 400 Hz and licking left in response to a 100 Hz standard. In a two-way ANOVA with an interaction of Frequency and Brain Region, performance in response to DCN stimulation was significantly better than for S1 (*p* = 0.015, F(1,24) = 6.8, two-way ANOVA). Dashed lines with hollow circles show d’ discrimination of hindlimb vibration trials (80% of trials), and solid lines and circles show performance from electrical microstimulation trials (20% of trials).

## Discussion

In the current study, we identified a circuit mechanism for the emergence of feature-selective encoding from peripheral mechanoreceptors along the ascending neuraxis, with a main transformation step at the level of thalamus. Traditionally, the somatosensory thalamus is viewed as an integration hub for sensory information from the periphery to higher brain regions (Shepherd and Yamawaki 2021; Halassa and Acsády 2016). In this study, we show that thalamus contributes to the transformation of neural coding from a temporal to a rate code. Specifically, we demonstrate for the first time that thalamic reticular nucleus plays a crucial role in transforming vibrotactile temporal coding into a rate code. The role of this inhibitory thalamic circuit is reminiscent of how cortical parvalbumin-positive interneurons improve feature selectivity, or how corticothalamic feedback sharpens spatial tuning in the mouse visual system (Born et al. 2021; S.-H. Lee et al. 2012). However, we cannot exclude contributions from other mechanisms of synaptic transmission (Bruno and Sakmann 2006). Future experiments, including cell patch clamp recording in brain slices of multiple regions along the ascending neuraxis could facilitate a more detailed characterization of the underlying synaptic mechanisms (Castro-Alamancos 2002; Mo et al. 2017; Yamawaki et al. 2021).

Our findings also suggest that inhibitory circuits within thalamus play a role in improving vibration frequency selectivity. In the whisker system of rats, topographic and stimulus-specific direct cortical excitation, and indirect inhibition through TRN, can selectively enhance or reduce spatial response tuning of ventral posteromedial nucleus neurons (Temereanca and Simons 2004), which is consistent with early evidence of lateral inhibition in the rat thalamus (Lavallée and Deschênes 2004; Pinault and Deschênes 1998).

A recent study (Turecek, Lehnert, and Ginty 2022) suggested that, in mice, a large proportion of high-frequency vibration information in the DCN is routed to the inferior colliculus but not to VPL, which largely contradicts previous work in multiple species (Harvey et al. 2013; Zhang et al. 2001; Herron and Dykes 1986). However, the authors used extracellular recordings along with antidromic activation to categorize projection targets of individual DCN neurons, which depend highly on the stimulus and recording electrode locations. We recently found that neurons responding to high frequency vibration are specifically arranged at the edge of the VPL and DCN (K.-S. Lee et al. 2023), which made it especially challenging to find thalamic projection neurons responding to high frequencies using antidromic activation and electrophysiological recording. Therefore, we performed retrograde viral labeling experiments targeting the thalamic VPL and the inferior colliculus to characterize the response properties of projection neurons, and discovered that both populations of projection neurons were encoding a broad range of frequencies (**Figure S9**).

Precise temporal coding in the somatosensory brainstem, at both the cellular and population level, might provide a significant advantage for potential brain machine interfaces (Loutit and Potas 2020a). Our behavioral experiments show, for the first time, that mice could discriminate brainstem electrical microstimulation frequencies. However, despite mice being able to discriminate high-frequency vibrations with very high accuracy, performance for discriminating high-frequency electrical microstimulation was relatively poorer. Indeed, mouse performance was comparable to that previously demonstrated by a macaque in a similar microstimulation task, only when the electrode quality was lower (Callier et al. 2020). This suggests higher quality electrodes that can be chronically implanted in the brainstem, are necessary to investigate the feasibility of a brainstem neural prosthetic device (Loutit and Potas 2020a). Nevertheless, Manipulation of temporal patterns of electrical pulses through microstimulation is readily achievable, and does not require characterization of detailed response properties, nor targeting of individual neurons (Stieger et al. 2022). In addition, the previously reported place code in DCN, where somatotopy and tonotopy were jointly observed, could be advantageous for reliably targeting neurons encoding spatial and temporal features of somatosensory stimuli, analogous to the artificial cochlea (K.-S. Lee et al. 2023).

A remaining challenge is to understand the perceptual relevance of vibration rate coding in thalamus and cortex. Because of the heterogeneity of thalamic response properties, the amount of convergence and divergence in downstream networks could complicate the effect of electrical stimulation. The hypothesis that activation of a frequency-tuned neuron directly links to perception of its preferred frequency could be directly tested by sensory mapping with two-photon calcium imaging combined with targeted multiphoton holographic optogenetics (Adesnik and Abdeladim 2021; Carrillo-Reid et al. 2019) or by using selective expression of channelrhodopsins such as a dual-protein switch system Cal-Light in the neural population responding to a targeted frequency (D. Lee et al. 2017). These experiments are technically still very challenging but might be necessary to unequivocally reveal the coding scheme.

Finally, we hope our findings will help to understand how feature selectivity emerges in sensory systems, as a transformation from temporal to rate coding is also common in the auditory and visual systems (Gao et al. 2016; Reinagel and Reid 2000). Moreover, detailed knowledge of the neural coding could assist the development of better neuroprosthetics in the ascending somatosensory pathway, and highlight the brainstem as a novel target for somatosensory microstimulation (Vickery et al. 2020; Loutit and Potas 2020a, 2020b).

## Legends

**Figure S1.**
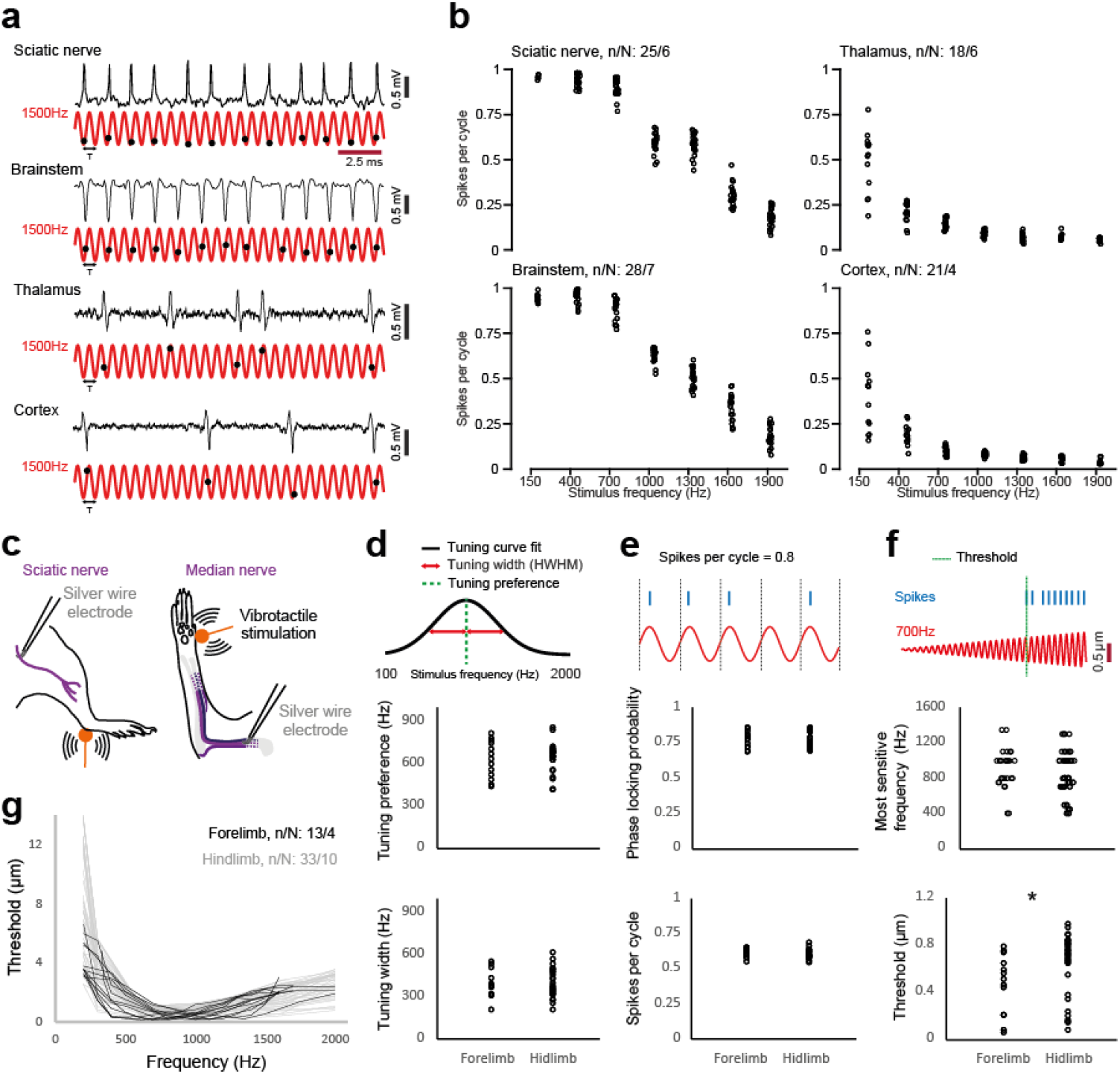
Response properties to forelimb and hindlimb vibration in the ascending pathway. **a,** Representative traces of four units spiking in response to 1500 Hz vibration stimulus. The timing of action potentials (black dots) relative to the periodic cycles of vibration (red) shows different levels of phase-locking. **b,** Spiking probability within individual cycles of vibration measured at each tested frequency, for nerve, brainstem, thalamus and cortex, respectively. **c,** Experimental setup for nerve fiber recording of sciatic nerve and median nerve. **d,** The distribution of the tuning preference (top) and tuning width (bottom) measured from recordings at the hindlimb and forelimb. **e,** The distribution of phase-locking probability (top) and spiking probability (bottom) measured from both the hindlimb and forelimb. **f,** The distribution of frequencies to which each unit was most sensitive frequency (top) and their thresholds (bottom) measured at both hindlimb and forelimb (unpaired t-tests, *P = 0.024). **g,** Frequency dependent threshold curves (mechanical activation threshold versus stimulation frequency) for both hindlimb and forelimb.

**Figure S2.**
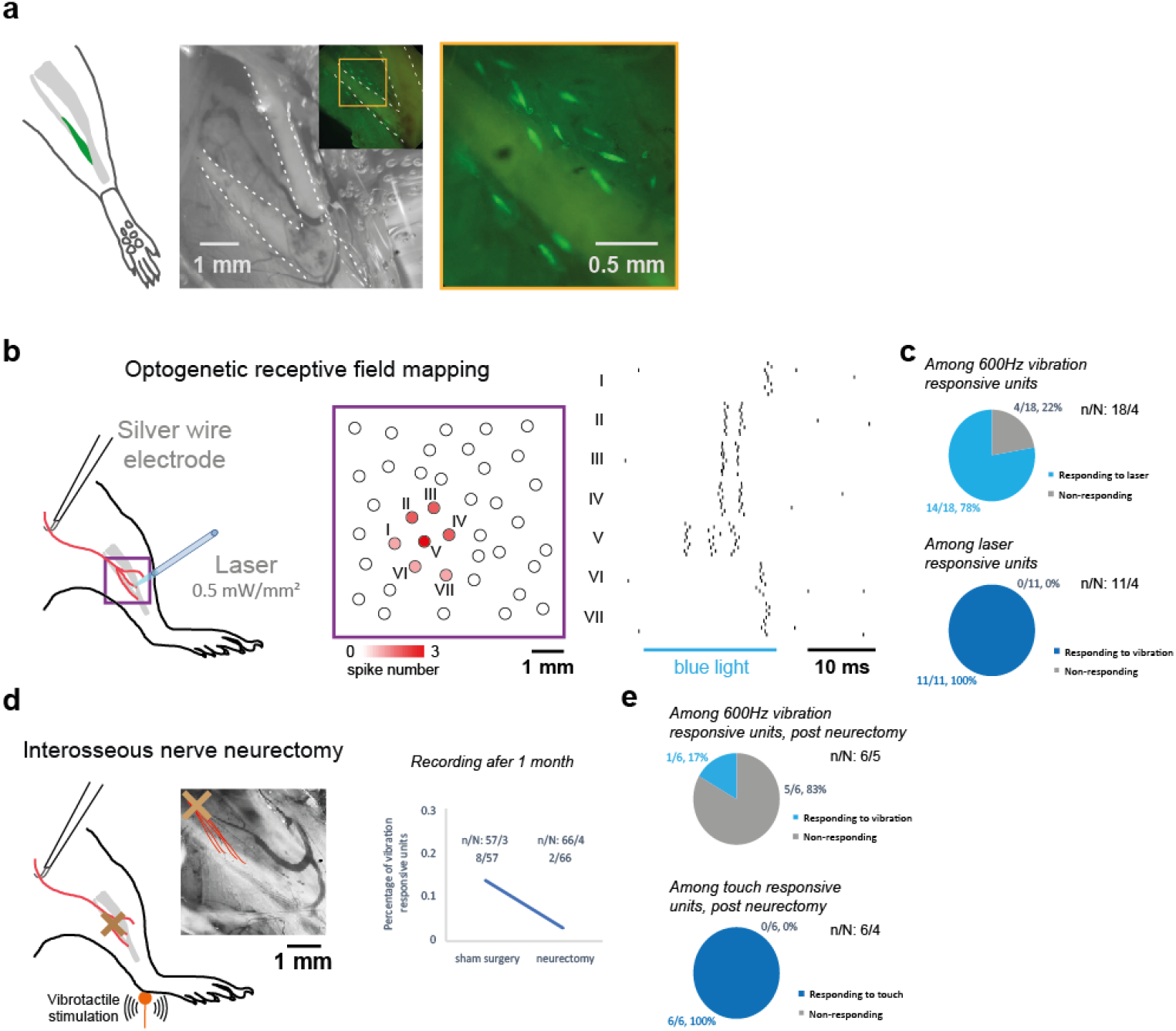
Identifying the origin of vibrotactile responses in the hindlimb. **a,** Distribution of Pacinian corpuscles on the fibula in a mouse hindlimb, identified by EYFP expression in the inner core structures (green). **b,** Left, optogenetic stimulation on Pacinian corpuscles in *Etv1-Cre* × *ChR2*-EYFP (Ai32) mice. Middle, blue light pulses with 0.5-1 mm diameter were manually directed to the exposed fibula area at each location indicated by a circle, without presumption of the location of Pacinian corpuscles. Right, an Aβ RA-LTMR innervated by a Pacinian corpuscle expressing *ChR2* in the inner core cells responded with action potentials when blue light was directed onto the location of the end organ. **c,** Top, out of 18 units that responded to 600 Hz vibration, 14 were activated by blue light illumination on the fibula. Bottom, all the laser responsive units are responding to vibration. **d,** Left, the schematic model of interosseous nerve neurectomy based on the nerve tracing. Right, after 1 month of neurectomy, terminal nerve recordings of vibration responsive units become very rare. **e,** During some terminal recording experiments, we directly compared the responsiveness of the same units pre- and post-neurectomy. Top, out of 6 units that responded to 600 Hz vibration, only 1 could still be activated by vibration post-neurectomy. Bottom, all the touch responsive units were not affected by neurectomy.

**Figure S3.**
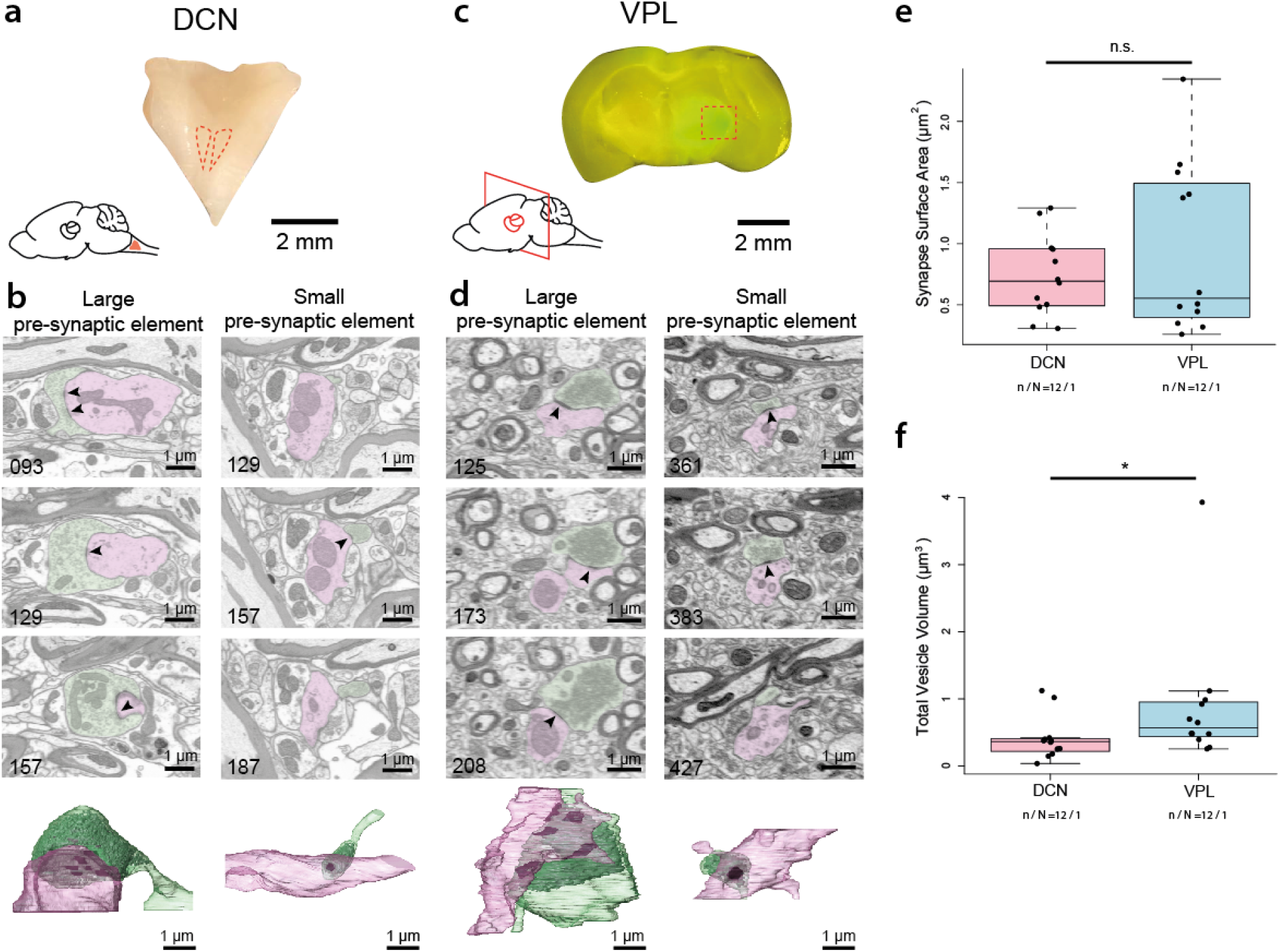
Electron microscopy analysis of synaptic structures of DCN and VPL. **a,** Dissected section of the brainstem. The dorsal column nuclei are indicated by the dotted red line. **b,** Two sample synapses in the DCN, one with a small and the other with a large presynaptic element. In the sample images from FIB-SEM, presynaptic elements are in green, postsynaptic elements are in pink and the arrowheads indicate synaptic electron-densities. In the respective reconstructions, presynaptic elements are in green, postsynaptic elements in pink, synapse in black and vesicles in gray. **c,** A GCaMP viral injection in hindlimb S1 retrogradely labels VPL projection neurons (green). This was used as a marker to identify VPL and dissect it for FIB-SEM. **d,** Same as b but the two sample synapses are from VPL. **e,** Distribution of total synaptic surface area per presynaptic element in DCN and VPL. Only synapses with large pre-synaptic elements (driver synapses) were included (n/N: 12/1, Wilcoxon rank-sum test, P = 0.7950). **f,** Distribution of total vesicle volume per presynaptic element in DCN and VPL (n/N: 12/1, Wilcoxon rank-sum test, P = 0.0304). Only synapses with large pre-synaptic elements (driver synapses) were included.

**Figure S4.**
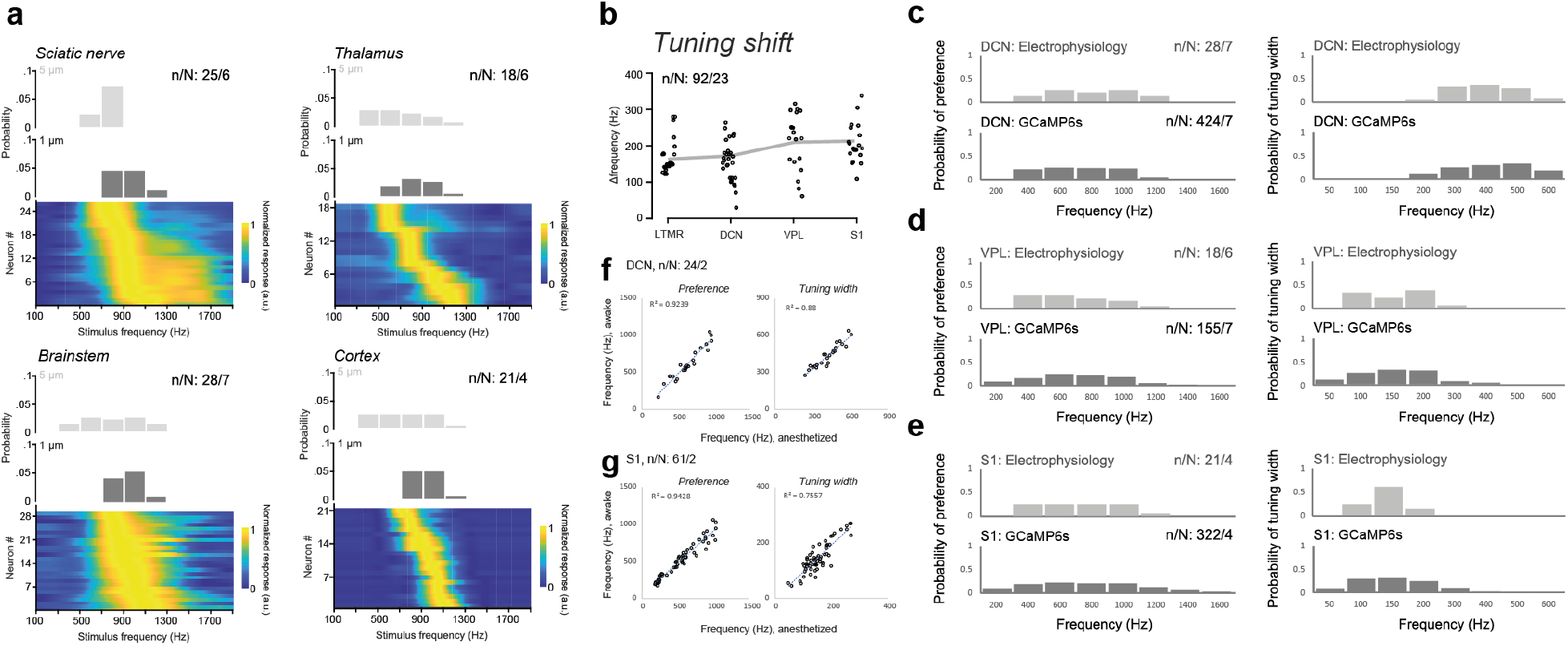
Frequency tuning depends on stimulus amplitudes, but not on recording methods. **a,** Normalized tuning curves of all neurons with identifiable tuning curve peaks sorted by decreasing best frequency and distribution of neurons’ best frequencies tested at 5 µm (top) and 1 µm (middle and bottom) amplitude. **b,** The tuning shift for the preferred vibration frequency in different vibration amplitudes, compared between all four areas in the ascending pathway (Kruskal–Wallis test, P = 0.567). **c,** Left, the distribution of tuning preferences measured from electrophysiology recordings (top) or calcium imaging (bottom) at DCN. Right, the distribution of tuning widths measured from electrophysiology recordings (top) or calcium imaging (bottom) at DCN. **d-e,** same as c, but for VPL and S1, respectively. No significant difference was found in any comparison between the results from electrophysiology and from imaging with Kolmogorov-Smirnov test. **f,** Left, the correlation of the frequency preference of individual DCN neurons measured with calcium imaging either under anesthesia or awake conditions. Right, the correlation of the tuning width of individual neurons (Pearson’s correlation, P < 0.0001). **g,** Left, the correlation of the frequency preference of individual S1 neurons measured with calcium imaging either under anesthesia or awake condition. Right, the correlation of the tuning width of individual neurons (Pearson’s correlation, P < 0.0001).

**Figure S5.**
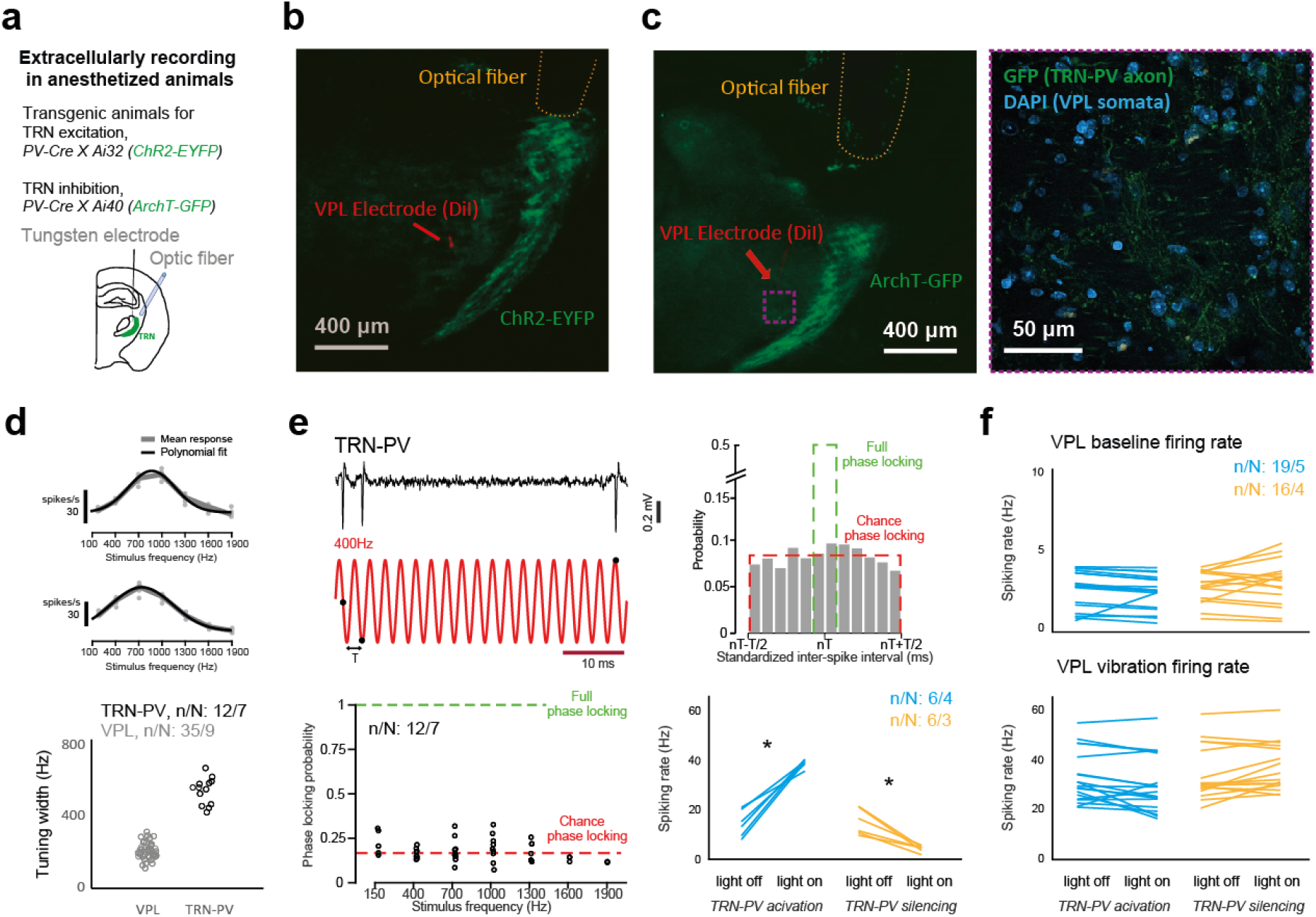
Properties of TRN-PV neuron responses to vibration and effects of optogenetic stimulation on thalamic nuclei. **a,** Experimental setup for paired optogenetic stimulation and extracellular recordings in thalamic nuclei. **b,** *Post hoc* histology demonstrates the YFP expression of TRN-PV neurons in *PV-Cre* X Ai32 mice and the electrode/optical fiber implantation can be localized. **c,** Left, coronal brain slice shows the GFP expression of TRN-PV neurons in *PV-Cre* X Ai40 mice and the electrode/optical fiber implantation can be localized. Right, the GFP signal within VPL is only present in axons, not in somas. **d,** Top, tuning curves based on mean spike rates of two opto-tagged TRN neurons. Bottom, the distribution of tuning widths of VPL and TRN-PV neurons (Wilcoxon rank-sum test, P < 0.0001). **e,** Top left, representative trace of a TRN-PV neuron spiking in response to 400 Hz vibration stimulus. The timing of its action potentials (black dots) is overlaid with periodic cycles of the vibration (red). Top right, distribution of standardized inter-spike intervals of an example neuron at this frequency. Bottom left, phase-locking probability measured at each tested frequency. Bottom right, the spiking rate of TRN-PV neurons affected by optogenetic activation or inactivation (paired-sample t-test, *P < 0.0001). **f,** Top, the baseline spiking rate of VPL neurons affected by optogenetic activation or inactivation of TRN-PV neurons (paired-sample t-test, P = 0.3478 and 0.1295). Bottom, spiking rate during a vibration stimulus of VPL neurons affected by optogenetic activation or inactivation of TRN-PV neurons (paired-sample t-test, P = 0.0324 and 0.0537).

**Figure S6.**
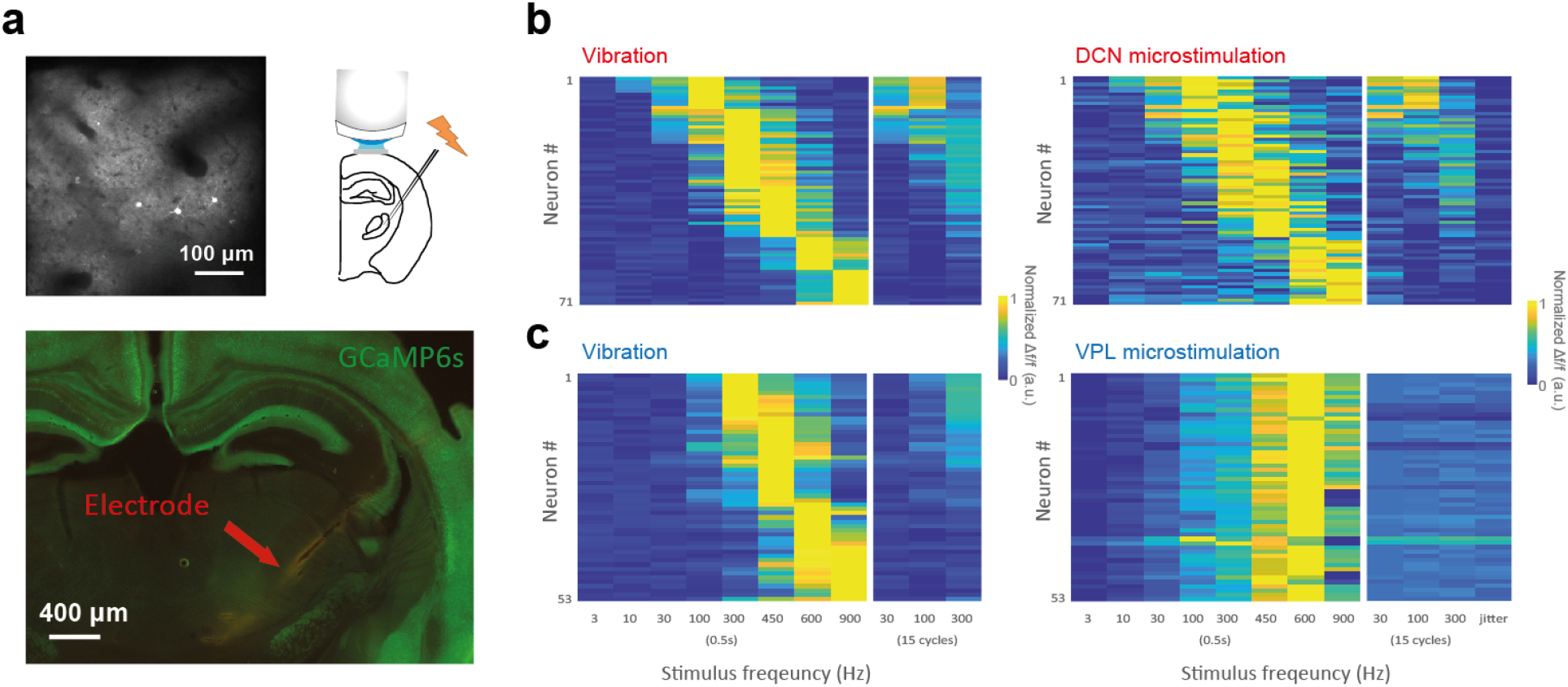
Microstimulation in DCN and VPL with readout in S1. **a,** Top, experimental setup for two-photon calcium imaging paired with electrical stimulation in VPL and an example field-of-view in S1. Bottom, an example tract of the electrode in histology. **b,** Left, frequency-selective tuning curves of S1 neurons in response to vibrations of different frequencies and durations. Right, frequency-selective tuning curves of the same S1 neurons in response to DCN microstimulation of different patterns and durations. c, Left, frequency-selective tuning curves of S1 neurons in response to vibrations of different frequencies and durations. Right, frequency-selective tuning curves of the same S1 neurons in response to VPL microstimulation of different patterns and durations.

**Figure S7.**
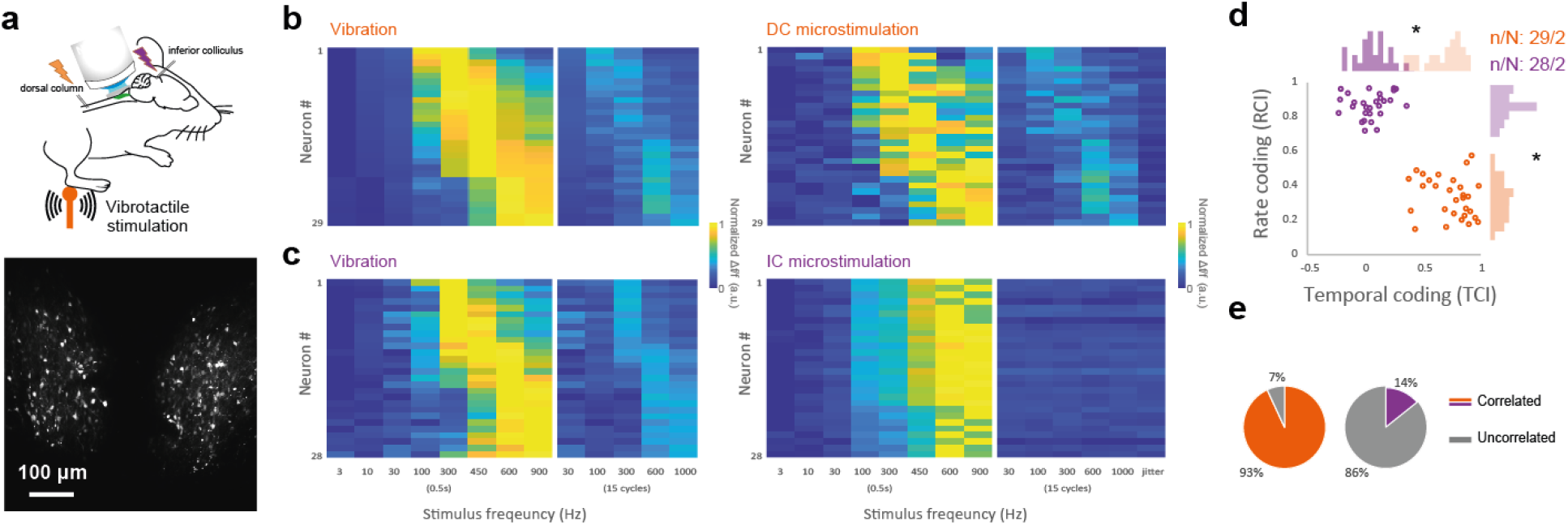
Microstimulation in dorsal column of lumbar spinal cord and inferior colliculus with readout in DCN. **a,** Top, experimental setup for two-photon calcium imaging paired with electrical stimulation in the dorsal column (DC) of lumbar spinal cord and inferior colliculus (IC), the unidirectional feedforward target of DCN. Bottom, an example field-of-view in DCN. **b,** Left, frequency-selective tuning curves of DCN neurons in response to vibrations of different frequencies and durations. Right, frequency-selective tuning curves of the same DCN neurons in response to DC microstimulation of different patterns and durations. **c,** Left, frequency-selective tuning curves of DCN neurons in response to vibrations of different frequencies and durations. Right, frequency-selective tuning curves of the same DCN neurons in response to IC microstimulation of different patterns and durations. **d,** The distribution of temporal coding index (TCI) and rate coding index (RCI) in both DC (orange) and IC (purple) showed very different coding strategies (unpaired t-tests, *P < 0.0001). **e,** The percentage of the DCN neurons whose vibration tuning curve can be reproduced by microstimulation tuning curves (Pearson’s correlation).

**Figure S8.**
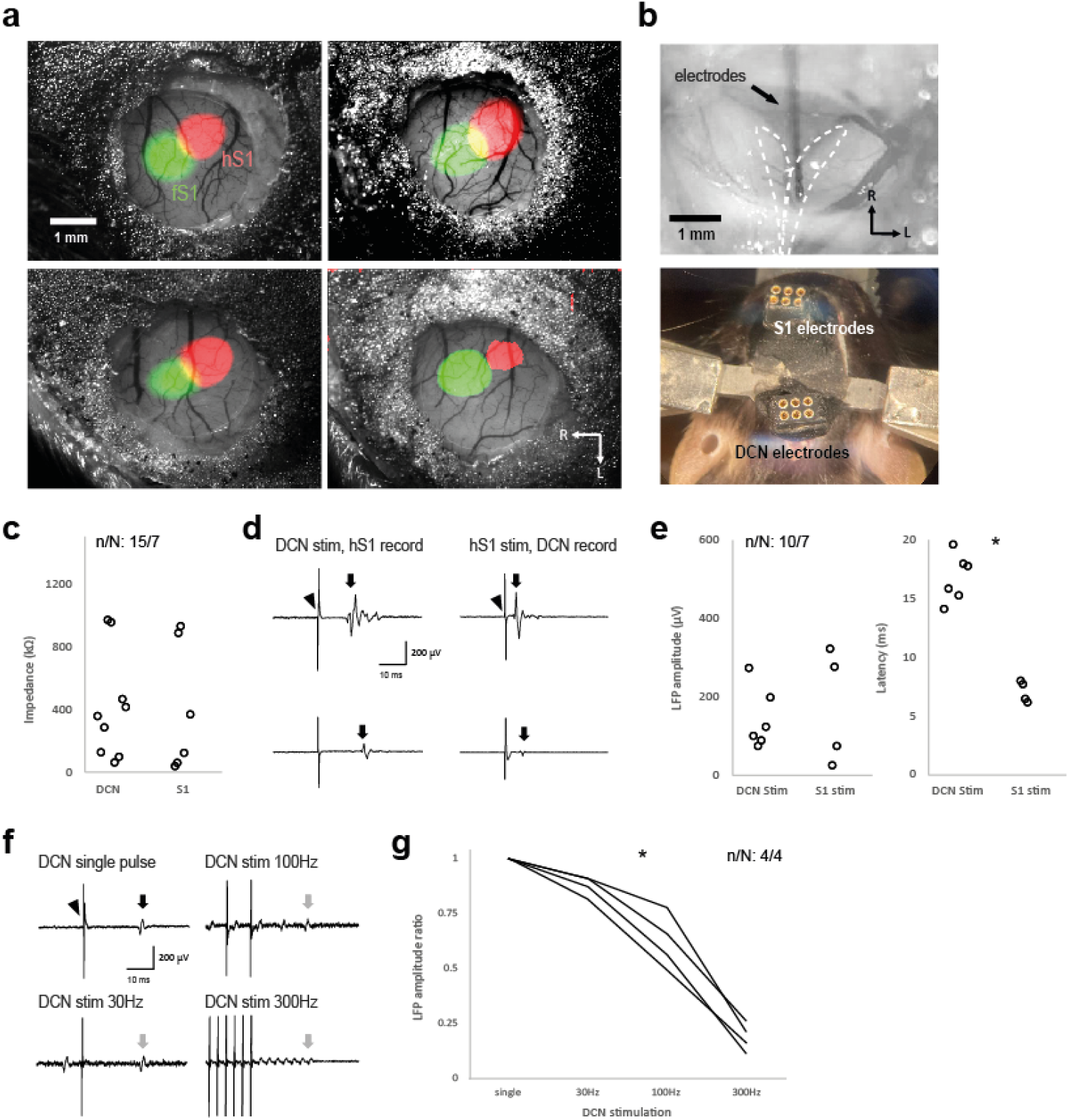
Validation of microstimulation with local field potential recording. **a,** An example of cranial windows and locations for electrode implantation. The targeted area is determined based on the position of the hindlimb S1 (hS1, red) defined with intrinsic signal imaging, neighboring to the forelimb S1 (fS1, green). **b,** Top, dorsal view of dorsal column nuclei (DCN) during electrode implantation. White dashed lines indicate the boundaries of the left and right gracile nuclei. Bottom, 5 weeks after the surgery. **c,** The impedance of electrodes in DCN and S1 after at least 3 weeks of implantation. **d,** Left, two example DCN stimulus-evoked local field potential (LFP) responses in hS1 (arrow). Right, two example hS1 stimulus-evoked local field potential responses in DCN (arrow). Arrowhead indicates the stimulus artifact. **e,** Left, the LFP amplitudes across all recordings. Right, the latency of LFP across all recordings (Wilcoxon rank-sum test, P < 0.0001). **f,** Example DCN stimulus-evoked LFP in hS1 with different stimulation patterns. Gray arrows indicate the LFP responding to the last electrical pulse of a 0.5 second stimulation train. g, Normalized LFP amplitudes across different stimulation frequencies (one-way repeated measures ANOVA, P < 0.0001). All LFP traces were averaged across 100 trials.

**Figure S9.**
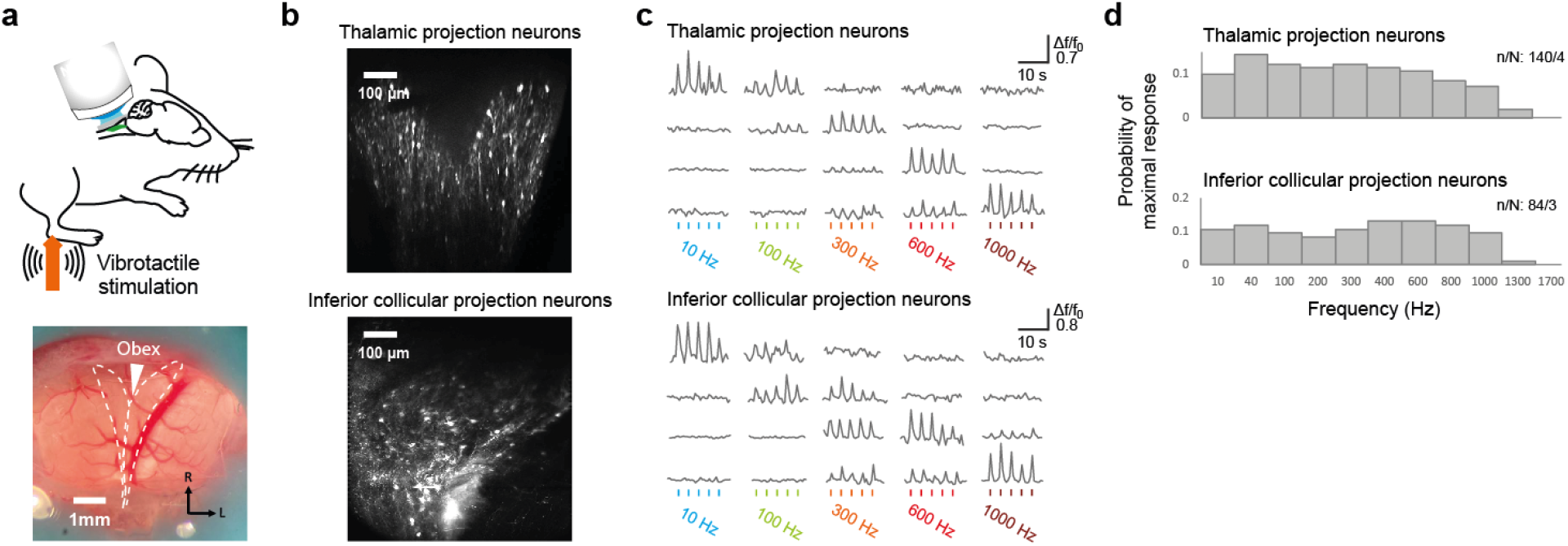
Response properties of thalamic and inferior colliculus projection neurons in DCN. **a,** Top, experimental setup schematic for two-photon calcium imaging. Bottom, dorsal view of the imaging window above the dorsal column nuclei (DCN). White dashed lines indicate the boundaries of the left and right gracile nuclei. The white arrowhead indicates obex and the midline. **b,** The retrograde expression patterns of GCaMP6s in DCN by injecting virus into the ventral posterolateral nucleus of the thalamus or inferior colliculus. **c,** Relative changes in fluorescence (Δf/f0) of eight example neurons imaged with two-photon microscopy in response to vibration of different frequencies. Coloured bars below indicate stimulus periods. **d,** The distribution of maximally-responding frequencies from individual neurons measured from calcium imaging of a population of thalamic projection neurons (top) or inferior colliculus projection neurons (bottom).

## STAR+METHODS

### KEY RESOURCES TABLE

| REAGENT or RESOURCE | SOURCE | IDENTIFIER |
| --- | --- | --- |
| Antibodies |  |  |
| Mice C57BL/6J | The Jackson Laboratory | JAX 000664 |
| Mice Ai32 | The Jackson Laboratory | JAX 012569 |
| Mice Ai40 | The Jackson Laboratory | JAX 021188 |
| Mice PV-Cre | The Jackson Laboratory | JAX 017320 |
| Mice CaMKII-tTA | The Jackson Laboratory | JAX 007004 |
| Mice TRE-GCaMP6s | The Jackson Laboratory | JAX 024742 |
| Mice ER81/Etv1 | The Jackson Laboratory | JAX 013048 |
| Bacterial and Virus Strains |  |  |
| AAV1.Syn.GCaMP6f.WPRE.SV40 | UPenn Vector Core | CS0939 |
| AAV1.Syn.GCaMP6s.WPRE.SV40 | UPenn Vector Core | CS0205 |
| AAVrg.Syn.jGCaMP7b.WPRE | Addgene | v63074 |
| AAV5.CAG.GCaMP6s.WPRE.SV40 | Addgene | v20751 |
| AAV1.CAG.GCaMP6s.WPRE.SV40 | UPenn Vector Core | V3096TI-3 |
| Software and Algorithms |  |  |
| Matlab | MathWorks | https://www.mathworks.com/ |
| Suite2p | https://github.com/cortex-lab/Suite2P | Pachitariu et al., 2016 |
| Fiji / ImageJ | https://imagej.net/software/fiji/ | Schindelin et al., 2012 |

### RESOURCE AVAILABILITY

#### Lead contact

Further information and requests for resources and reagents should be directed to and will be fulfilled by the lead contact, Kuo-Sheng Lee or Daniel Huber.

#### Materials availability

This study did not generate new unique reagents.

#### Data and code availability

All data reported in this study and code used for analysis will be shared by the lead contact upon request.

### EXPERIMENTAL MODEL AND SUBJECT DETAILS

Electrophysiology and imaging experiments at different parts of the ascending pathway were conducted with C57BL/6 (Charles River Laboratory) mice. Experiments involving optogenetic manipulation of thalamic inhibitory circuits were conducted in double-transgenic mice generated by mating homozygous Ai32 mice carrying a floxed *ChR2*-EYFP fusion gene inserted in the *Gt(ROSA)26Sor* locus in a C57BL/6 strain (Jackson Laboratory; stock no. 012569) or homozygous Ai40 mice carrying a floxed *ArchT*-EYFP fusion gene inserted in the *Gt(ROSA)26Sor* locus in a C57BL/6 strain (Jackson Laboratory; stock no. 021188) with homozygous PV-Cre (Jackson laboratory; stock no. 017320) mice. As such, Cre-mediated recombination resulted in an expression of the floxed opsin sequence in the PV-expressing cells of the offspring. Imaging experiments involving calcium imaging in S1 were conducted in double-transgenic CaMKII-GCaMP6s mice generated by mating heterozygous CaMKII-tTA mice (Jackson Laboratory; stock no. 007004) and TRE-GCaMP6s mice (Jackson Laboratory; stock no. 024742). CaMKII-GCaMP6s mice express the neural activity indicator GCaMP6s (T.-W. Chen et al. 2013) in excitatory neurons of the cortex (Wekselblatt et al. 2016). Hindpaw optogenetic stimulation of Pacinian corpuscles was conducted in double-transgenic mice generated by mating homozygous Ai32 mice with heterozygous ER81/Etv1-CreER mice expressing the CreER^T2^ fusion protein from the *ER81/Etv1* promoter elements (Jackson Laboratory; stock no. 013048). As such, Cre-mediated recombination resulted in an expression of the floxed ChR2 sequence in the ER81/Etv1-expressing cells of the offspring. ER81/Etv1 is expressed in the inner core region of Pacinian corpuscles. To induce CreER-based recombination, double-transgenic adult offspring were administered one daily dose of a 100 µL tamoxifen (T5648, Sigma-Aldrich) solution (20 mg/mL) dissolved in corn oil (C8267, Sigma-Aldrich) for five consecutive days by intraperitoneal injection. Wild type mice were purchased from Charles River and transgenic mice were obtained from The Jackson Laboratory and bred in the animal facility of the University of Geneva.

Mice were housed in an animal facility, maintained on a 12:12 light/dark cycle, and the experiments were performed during the light phase of the cycle. The animals did not undergo any previous surgery, drug administration or experiments and were housed in groups of maximum 5 animals per cage. All procedures complied with and were approved by the Institutional Animal Care and Use Committee of the University of Geneva and Geneva veterinary offices.

### METHODS DETAILS

#### Surgery

15 to 30 week old mice were surgically prepared for terminal two-photon Ca^2+^ imaging. Surgeries were conducted under isoflurane anesthesia (1.5 to 2%) and additional analgesic (0.1 mg/kg buprenorphine intramuscular (i.m.)), local anesthetic (75 µL 1% lidocaine subcutaneous (s.c.) under the location for incision) and anti-inflammatory drugs (2.5 mg/kg dexamethasone i.m. and 5 mg/kg carprofen s.c.) were administered. Mice were fixed on a bite bar with a snout clamp and rested on top of a heating pad. For viral expression at the brainstem, thalamus and cortex, the scalp was cut over the midline between the ears and eyes. After removing the dura, a bulk viral injection (400 to 800 nL) was administered with pulled and beveled glass pipettes (≈15 µm tip diameter) at an approximate rate of 20 nL/min into either, the gracile nucleus right next to the obex, the ventral posterolateral nucleus (VPL) of the thalamus (2 mm lateral and 2 mm posterior to bregma; 3.5 mm below the dura), inferior colliculus (2 mm lateral and 0.5 mm posterior to lambda; 350 µm below the dura), or the hindlimb representation of the primary somatosensory cortex (2 mm lateral and 0 mm anterior to bregma; 350 µm below the dura). For a dense expression among the neural population, the injections consisted of AAV1.Syn.GCaMP6f.WPRE.SV40 (University of Pennsylvania, CS0939, titer 6.93 x 10^13^), AAV1.Syn.GCaMP6s.WPRE.SV40 (University of Pennsylvania, CS0205, titer 3.22 x 10^13^), or AAVrg.Syn.jGCaMP7b.WPRE (Addgene, lot. no. v63074, titer 1 x 10^13^). For GCaMP expression in the DRG neurons, AAV5.CAG.GCaMP6s.WPRE.SV40 (Addgene, lot. no. v20751, titer 3.2 x x 10^13^), or AAV1.CAG.GCaMP6s.WPRE.SV40 (University of Pennsylvania, V3096TI-3, titer 2.33 x 10^13^) was directly injected (1000 to 3000 nL) into the lumbar enlargement in between T13 and L4 vertebrates. During the initial development of DRG and DCN imaging, we tested different GCaMP variants and AAV serotypes. In the end, the physiological results were not significantly different between different AAV serotypes and GCaMP variants (K.-S. Lee et al. 2023). For visualizing the process of injection, 0.1 μL of 2% FastGreen was added into 5 μL of the virus. After the end of each injection, the pipette was left in place for at least another 10 min before being retracted. After 3-8 weeks of expression and recovery, experiments could be conducted.

All imaging and electrophysiology experiments were surgically prepared under terminal anesthesia by inhalation of isoflurane (∼2%) and the body temperature was maintained near 37 °C. For analgesia, buprenorphine (0.1mg/kg SC) was provided 15-20 min before the procedure, except for thalamus and cortex, where chronic preparation was employed. An additional dose of buprenorphine was given if the procedure exceeded 4 hours. The whole procedure was limited to 8 hours. For imaging of brain structures, a custom-made titanium head bar was fixed on the skull with a cyanoacrylate adhesive (ergo 5011, IBZ Industrie) and dental cement to allow subsequent head fixation. After the initial surgical phase, the animal was maintained under light isoflurane anesthesia (0.25 – 0.75%) and allowed at least 20 minutes of recovery before recording.

Imaging of dorsal root ganglia (DRG) was done at least four weeks after GCaMP expression. The surgical procedure for vertebral window implantation was adapted from a previous study (C. Chen et al. 2019). In short, after hemostasis, DRG was covered with a thin layer of Kwik-Sil silicone elastomer (World Precision Instruments) and sealed with a 2-mm diameter cover glass, and the cover glass was attached to the vertebral mount by cyanoacrylate glue and dental acrylic.

For brainstem imaging, the skin and cervical paraspinal muscle over the neck were cut and pulled to the side. A custom made titanium head bar was fixed on the skull with a cyanoacrylate adhesive (ergo 5011, IBZ Industrie) and dental cement. The head was fixed at approximately 45° downward. After the removal of dura and the skull above the posterior part of the cerebellum, Kwik-Sil was applied to the surface of the brainstem before a hand-cut wedge-shaped glass coverslip (150 µm thickness) was placed over the brainstem. The glass window over the gracile nucleus was first stabilized by the pressure of a micromanipulator. The micromanipulator was removed after the cyanoacrylate glue and dental acrylic hardened. Due to the fact that the DCN of those animals with viral injection in thalamus or inferior colliculus will show neurons retrogradely labeled from their axonal terminals in these target areas, we performed DCN imaging in those animals to investigate the tuning properties of thalamic projection neurons and inferior colliculus projection neurons.

For thalamic and cortical imaging, survival surgery for window implantation was performed either right after viral injection or after 2-4 weeks of recovery period and viral expression. A craniotomy was performed over the target structures. For cortical imaging, two hand-cut glass coverslips (150 µm thickness) matched to the shape of the craniotomy were glued together with optical adhesive (NOA 61, Norland Products).The lower glass was placed on top of the cortex, and the upper glass, which was larger than the craniotomy, was glued to the skull with cyanoacrylate adhesive and secured with dental cement. For the cortical imaging using transgenic mice, viral injections were skipped. For thalamic imaging, a gradient-index (GRIN) lens (Inscopix) was used instead of a glass coverslip. The correspondence between the injection sites and the sensory hindpaw representation was confirmed during *in vivo* sensory mapping.

#### Two-photon Ca^2+^ imaging

Imaging was performed with a custom-built two-photon microscope (MIMMS, www.openwiki.janelia.org) controlled by Scanimage 5.1 (Vidrio Technologies) using a 16X 0.8 NA objective (Nikon) and with excitation wavelength set to ∼920 nm (Ultra II, tunable Ti:Sapphire laser, Coherent). 512 by 512 pixel images were acquired at 29.72 Hz using bidirectional scanning with a resonant scanner system (Thorlabs). The power was modulated with a Pockels cell (350-80-LA-02, Conoptics) and calibrated with a photodiode (Thorlabs). The primary mirror for imaging was a custom polychroic (Chroma, zt470/561/nir-trans) that could transmit infrared light, while reflecting green light. Before detection, the remaining infrared light was filtered with a coloured glass band pass filter (BG39, Chroma). Images were acquired using photomultiplier tubes (H11706P-40 SEL, Hamamatsu) and written in 16 bit format to disk.

Raw calcium images were processed using Suite2P, a publicly available two-photon calcium imaging analysis pipeline (Pachitariu, Stringer, and Harris 2018). First, images were registered based on the cross-correlation of the averaged images to account for brain motion. Then, regions of interest (ROI) were established by clustering neighboring pixels with similar time courses. Next, manual selection of ROI was performed to eliminate low-quality or non-targeted regions of interest.

#### Intrinsic signal imaging

Intrinsic signal imaging was performed to precisely localize the hindlimb representation of S1 for electrode implantation. Mice were head-fixed and placed on a heated platform under light isoflurane anesthesia (0.25 to 0.75%). Their right forepaw or hindpaw rested on a metal rod (2 mm diameter) mounted on a galvanometric actuator (model. no. 000-G120D, GSI Lumonics). Fifteen 1-s vibrotactile stimuli consisting of a 100 Hz sinusoidal vibration were delivered to the forepaw with a 10 s inter-stimulus interval. The cranial window was illuminated with a collimated red light LED (635 nm peak, ACULED VHL, part. no. E001947, Perkin Elmer) and imaged with a CCD camera at 10 fps with 256 × 332 pixel resolution (Retiga-2000R, Q Imaging). The average difference image between the stimulation period and a 1.5-s baseline period was processed using a 50 × 50-pixel spatial averaging filter and subsequently smoothed with a 5 × 5-pixel Gaussian lowpass filter with 0.5 pixel s.d. Stimulus presentation and image acquisition was controlled using Ephus software (https://Scanimage.org).

#### Electrophysiology

For electrophysiology experiments, mice were anesthetized by isoflurane inhalation (2%) and body temperature was maintained near 37°C with a heating mat. For analgesia, buprenorphine (0.1mg/kg SC) was provided 15-20 min before the procedure. To assess the sensory afferents innervating Pacinian corpuscles, a skin opening was performed along the hamstring muscle. Then, the sciatic nerve was carefully isolated using ophthalmic scissors and tweezers. The pia mater spinalis and dura mater around the nerve were removed, and finally, the nerve bundle was separated into single fibers (15-20 μm in diameter) and placed on two Ag/AgCl wire electrodes in the recording pool filled with mineral oil. A third ground Ag/AgCl electrode was placed in the muscle next to the recording chamber.

For all recordings in the central nervous system, a tungsten monopolar microelectrode (FHC, 30031) was used. A small craniotomy was drilled for S1 and VPL recordings in mice surgically implanted with a head bar as described above. We sampled neurons from cortical layer 2/3 (above 400 μm) as it is more comparable to our calcium imaging from cortical layer 2/3. Though the major thalamocortical inputs from VPL first arrive at layer 4, our previous results recorded from all cortical layers observed no difference in coding for vibration among the dataset (Prsa et al. 2019).

To characterize the response properties of sensory afferents at the peripheral level, we recorded stimulus-evoked activity of afferents *in vivo* using hook electrodes in the both tibial nerve and median nerve. Vibratory stimuli were applied to different locations of the hindlimb or forelimb using a calibrated piezo stimulator (Physik Instrumente P-841, E509 controller, E504 amplifier), so that the neural responses to all combinations of locations, frequencies and amplitudes could be characterized. To determine the mechanical sensitivity threshold and frequency tuning of each afferent, a series of sinusoidal mechanical stimuli (duration 20 s for frequencies above 100 Hz; linearly increasing amplitude) were applied to the hand-mapped receptive field of individual afferents with the piezo actuator.

The signal was amplified and filtered (>10 Hz and <10 kHz) and acquired at 30 kHz (PXIe-1073, National Instruments) using WaveSurfer (https://wavesurfer.janelia.org/) Matlab (Mathworks) routines. Trial start, stimulus onset triggers, and details of stimulus were saved in parallel on separate channels and used for *post-hoc* alignment of recorded spikes. After the recording, animals were euthanized by overdose with isoflurane (5%) followed by cervical dislocation and bleeding.

#### Vibrotactile stimuli

The vibrotactile stimuli were generated by a bimorph piezoelectric multilayer bender actuator with a piezoelectric stack actuator (P-841.3, Physik Instrumente) equipped with a strain gauge feedback sensor for high frequency stimulations (above 100 Hz). A hand-cut blunt plastic cone was mounted on the actuator. The actuator and sensor controllers (E-662, E-618.1 and E-509.S1, Physik Instrumente) operated in an open loop with a contacting force below 50 mN on the mouse skin. The forces of our vibrotactile stimuli measured at the static states of 3µm, and 30 µm, were less than 10 mN, and 10-30 mN. Pure sinusoids (250 or 500 ms duration) of a wide range of frequencies (100 to 1900 Hz) calibrated to produce a desired displacement amplitude (0.01 to 20 µm) were sampled at 10 kHz (USB-6353, National Instruments) and fed to the actuator controller. The sensor measurements were continuously acquired and recalibration of motor commands was regularly performed for the stimuli to remain highly consistent, by comparing the ground truth data acquired optically with a laser doppler vibrometer system (Polytec, OFV-5000). The spectrum of the acquired sensor measurements was analyzed to ensure the integrity of their frequency content (Prsa et al. 2019; K.-S. Lee et al. 2023).

#### Optogenetic stimuli

Optogenetic stimulation of parvalbumin neurons in thalamic reticular nucleus (TRN-PV) was generated by blue light illumination (473 nm, 50 mW; Obis LX FP 473, Coherent) out of a fiber (3.5 µm core diameter, 0.045 NA) operated in analog control mode or by yellow light illumination (568 nm, 500 mW; Sapphire LP/LPX, Coherent) fed into an optic fiber with achromatic FiberPorts (Thorlabs), gated with a shutter. The laser power or the shutter was controlled by analog signals from the WaveSurfer (https://wavesurfer.janelia.org/). For the blue laser, the stimulus was a 40-Hz train of square pulses (2000 ms duration, 1 ms pulse length). For the yellow laser, the stimulus was a continuous square pulses (2000 ms duration). The mean power of the stimulation signal used in our experiments was 1.2 mW for blue and 0.5 mW for yellow (measured at the tip of the fiber).

Fiber-optic implantation in ipsilateral TRN involved cutting an optic fiber (high OH, 400 µm core, 0.39 NA, Thorlabs) with a diamond knife followed by insertion into the 1.25 mm diameter of a ceramic ferrule (Thorlabs) with a 440 µm bore (K.-S. Lee et al. 2019). The optic fiber was adhered to the ferrule with epoxy (Gorilla Glue Company). Both ends of the optic fiber were finely ground with sandpaper and grinding puck (Thorlabs). The optic fiber was implanted into the thalamic reticular nucleus with a 45° angle laterally. Dental cement (C&B Metabond) was applied to adhere the optic fiber to the surface of the skull. For optical stimulation of Pacinian corpuscles, the stimulus was 20 ms short square pulses with near threshold intensity. Stimulation of different locations on the hindpaw was achieved by manually directing the optical fiber to a 7 × 7 mm area, and the laser beam was focused with FiberPorts (Thorlabs), which focused the light to a 474-μm diameter spot (measured with a beam profiler (Laser Beam profiler Spiricon, SP620U and BeamMic software).

#### Electrical stimuli

For terminal microstimulation experiments in both S1 and DCN, a craniotomy above the target region was performed and bipolar electrodes (platinum–iridium 250 µm spacing, FHC) were lowered into the contralateral gracile nucleus, or the ipsilateral VPL of the thalamus (coordinates: electrode angled 45°, 2 mm posterior to bregma, 5.5 mm lateral, 5 mm deep) for S1 imaging. For DCN imaging experiments, electrodes were targeted at ipsilateral dorsal column of lumbar 4 to lumbar 6 spinal cord or the contralateral inferior colliculus (coordinates: 1.5 mm posterior to lambda, at tissue entry, 1.5 mm lateral, 1 mm deep).

For behavioral experiments, the stimulus electrodes were composed of two thin platinum–iridium wires (101R-1W, Science Products) twisted together and soldered directly to the connector. On average electrodes had an impedance of ∼1 kHz to <1000 kΩ, measured by a NanoZ impedance meter (Plexon). With intrinsic imaging data or the known somatotopy, one or two pairs of electrodes were implanted to the center of the hindlimb representation of left S1 (200–300 μm) or right posterior gracile nucleus (angled 45° rostrally, 100–200 μm), and the brain was covered by first Kwik-Sil silicone elastomer (World Precision Instruments), cyanoacrylate glue and dental acrylic.

The stimulus was generally a train of square pulses (500 ms duration, 0.2 ms pulse length) with various frequencies (50 to 600 Hz) and amplitudes (100–300 µA). Validation of microstimulation was also tested by stimulus evoked responses in calcium imaging in S1 or by local field potential recording in the projected areas after weeks of implantation and stimulation. For example, electrical stimulation in the gracile nucleus was verified by recording the stimulus evoked field potential from contralateral S1.

#### Neurectomy

The surgery was performed in mice under isoflurane anesthesia (1.5–2%) and additional analgesic (0.1 mg/kg buprenorphine i.m.), local anesthetic (30 µl 1% lidocaine s.c. in the hindlimb) and anti-inflammatory drugs (2.5 mg/kg dexamethasone i.m. and 5 mg/kg carprofen s.c.) were administered. The hair of the forelimb was shaved and a skin incision was made behind the knee. The muscles and connective tissues were retracted to expose the tibial nerves innervating the interosseous region. Complete transection of the nerve branch innervating the group of Pacinian corpuscles on the fibula was performed. After suturing the skin, mice were allowed to recover for 3 days before any follow-up experiments. The electrophysiological experiment was performed 1 month after surgery and the animals were euthanized. During some terminal nerve fiber recording experiments, the same procedure was performed at the end of the recording, to observe the immediate effects of neurectomy on different fiber types responding to vibration or touch.

#### Histology

Upon completion of terminal experiments, isoflurane was raised to 5% and 0.2 ml pentobarbital was delivered through an intraperitoneal injection. The animal was transcardially perfused with 20 ml of 0.9% NaCl and then 50 ml of 4% paraformaldehyde in 0.1 M PB. The brain or body parts were removed and placed in 4% PFA overnight. A vibratome (Leica VT1200S) was used to cut tangential or coronal sections (100 µm thick). Slices were washed three times using 1X PBS, then mounted to a slide using SlowFade Gold (Thermofisher Scientific) or Mounting Medium With DAPI (Ancam,104139) for imaging using a fluorescence microscope (Olympus BX53) or confocal microscopy (ZEISS LSM 800). In most of the recording in the deep brain areas, the electrode was coated with DiI (ThermoFisher) before insertion, for identification of the precise recording site.

#### Confocal imaging

Dissection of tissue was done under a fluorescent microscope (Leica M165FC). The expression pattern of GCaMP, ArchT or ChR2 was imaged by illuminating the dissected tissue with a blue LED light, with emission channels optimized for each fluorophore (GFP, or YFP), and subsequently imaged with confocal or two-photon microscopes. Confocal imaging was done using a ZEISS LSM 800 confocal microscope. Images were acquired using 405, 488, and 561 nm laser lines with emission channels optimized for each fluorophore (DAPI, GFP, DiI) while minimizing cross-talk between channels. Images were acquired at 512×512 with sampling resolution ranging from 0.15 to 1 μm per pixel. Z-stacks were acquired using 0.5 to 5 μm steps and the confocal pinhole was set to 1 AU. Colocalization was performed manually. The imaging data was analyzed using ImageJ (NIH).

#### Data analysis

##### Significant responses

For calcium imaging, ROIs were circular or drawn manually for the targeted neuronal structures in ImageJ (NIH). Fluorescence time-courses were computed as the mean of all pixels within the ROI at each time point, then extracted using Miji (Sage et al., n.d.), and finally synchronized with the stimulation. Evoked responses were computed as changes in fluorescence relative to baseline fluorescence. Frequency selectivity for individual neurons or nerve fibers was quantified by a previously reported method (K.-S. Lee, Huang, and Fitzpatrick 2016). To computing tuning properties, the fluorescence signal was calculated as Δf/f=(f–f0)/f0, where f0 is the baseline fluorescence signal averaged over the 1 s period immediately before the start of the vibration stimulation and f is the fluorescence signal averaged over the first 1.5 s period after the start of the stimulation. Tuning curves were obtained by calculating the mean fluorescence signal (Δf/f) for each stimulation, and then fitting a polynomial curve to the resulting data. For electrophysiological recording, tuning curves were computed by the spike rate of the neurons. To ensure that we were recording from a single nerve fiber/neuron and to avoid the complications of spike sorting, we only sampled units with a signal-to-noise ratio (SNR) > 5, when responding to a vibration stimulation (100 or 700 Hz). SNRs were calculated as the maximum amplitude of the mean spike waveform divided by the standard deviation of the background noise. There was no additional offline filtering performed for spike detection.

##### Tuning curve fitting

Neuronal structures were considered to be responsive if the maximum stimulus-related fluorescence response (Δf/f) or spiking responses to any stimulus was greater than 5% on average, and also greater than 2 standard deviations (SD) above the mean baseline fluorescence. In addition, we required that neurons responded at least 2 SD above baseline on at least 20% of the trials tested. The same criteria were used to identify neurons with significant responses to the vibration, optogenetic and electrical stimuli.

Neuronal structures were considered to be frequency selective if they were responsive and also met the following criteria: (1) well fit by the polynomial function (r > 0.7, p < 0.05), and (2) tuning index (TI) > 0.2

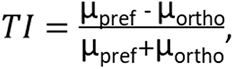

where μ_pref_ denotes the mean response to the preferred stimulus frequency and μ_ortho_ is the mean response to the least preferred stimulus frequency, defined by the curve fitting. To characterize a neuron’s tuning to vibration frequency, the normalized mean responses were fit to frequency by the polynomial curve fitting function (Matlab) using the method of non-linear least squares, with the degree of polynomial fit set at 6. The tuning preference of individual neurons was defined by the peak of the curve and the tuning width was defined by the half-width at the half-maximum of the curve (HWHM).

##### Phase-locked spiking

To determine whether a nerve fiber or a neuron was entrained by the sinusoidal vibration, we calculated standardized inter-spike intervals. We chose this measure because it is immune to variability in response onset times across different stimulus repetitions. For a given vibration frequency, the inter-spike intervals (ISI) of all possible spike pairs that occurred during stimulation were calculated and grouped across stimulus repetitions. Because entrained spiking should yield an ISI distribution that peaks at integer multiples of the sinusoidal stimulus period T, values were converted to standardized inter-spike intervals (SISI) according to SISI = T + (ISI − nT), where nT is the integer multiple of T closest to ISI. Entrainment probability was defined as the percentage of ISIs in the [nT − T/12, nT + T/12] interval. Given that standardized ISIs are distributed between nT − T/2 and nT + T/2, entrainment probability should equal unity in the case of perfect entrainment and be equal to 1/6 in the case of chance entrainment. For each neuron and at each frequency we repeated the calculation of entrainment probability 1,999 times with randomly sampled ISIs (with replacement). We then measured whether the lower 99th percentile of the repeated measures was less than 1/6, which constitutes a one-tailed bootstrap test at significance level P < 0.01. For complementary analysis of spiking regularity, we computed the coefficient of variation (CV), which is the standard deviation of the inter-spike intervals (ISI) divided by the mean ISI.

##### Temporal and rate coding index

Temporal coding index (TCI) and rate coding index (RCI) were computed for individual imaged neurons in S1 or in DCN by comparing their tuning curves to vibrotactile stimuli and electrical microstimulation of downstream brain areas. Temporal coding index (TCI) was defined as the reproducibility of the vibration-induced frequency tuning curve by temporally structured microstimulation, which mimics the spiking of the sensory afferents responding to vibration on the skin. TCI was defined as the correlation coefficient between tuning curves to vibration vs. microstimulation. On the other hand, rate coding index (RCI) was defined as the invariant responses from the microstimulation with various frequencies but with the same number of electrical pulses, which mimics the fixed number of spikes of the presynaptic inputs. In the case where the temporal precision is not critical and the response to the stimulation is only determined by the number of spikes, the neural responses of the postsynaptic neuron should be the same for all stimulation with different temporal patterns under the control of the inputs’ spiking rate. We considered the whole calcium trace following the stimulation as a response from the cortical neuron, as calcium imaging is empirically sensitive to the time scale in this range.

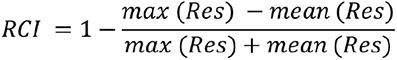

where Res denotes the responses to all four microstimulation patterns with different frequencies (30, 100, 300 Hz with equal periods, and 100 Hz jittered by ±3 ms, from a uniform distribution) but with the same number of 15 pulses. In order to test the robustness of the RCI index, we also computed the correlation between the RCI with RCI_same length_, which only included the response to the regular and jittered 100 Hz pulse trains. The results were highly consistent for both DCN and VPL stimulation (R^2^: 0.7163 and 0.4164).

#### Statistics

No statistical methods were used to predetermine sample size and all experimental animals were included in the analysis. The normality assumption was tested with the Kolmogorov-Smirnov test. Non-parametric tests were used when the normality assumption was not met. We used a two sided non-parametric Wilcoxon rank-sum test to compare two groups and the Kruskal–Wallis test to compare multiple groups with *post-hoc* tests using Dunn’s test, without assumptions of normality or equal variances. Paired-sample t-test was used to compare the effect of optogenetic manipulations. Pearson’s correlation coefficient was applied to the analysis related to frequency tuning. All statistical methods were two-sided. All data analyses were performed with custom written routines in Matlab.

#### Behavioral task

##### Microstimulation frequency discrimination

Mice were prepared for microstimulation frequency discrimination tasks by fixing a custom-made titanium head bar to the skull with a cyanoacrylate adhesive (ergo 5011, IBZ Industrie) and dental cement, and implanting the stimulating electrodes in the hindlimb region of the DCN and S1, under general isoflurane anesthesia (1.5–2%). After surgical recovery, mice were placed under a water restriction regime (1ml/day) and habituated to the tunnel and head-fixation frame where they performed the task.

Mice were trained daily to discriminate vibration or electrical microstimulation at lower frequencies from stimuli at higher frequencies in two-alternative forced-choice (2AFC) tasks. Each trial began with a 1-s hold period in which mice had to refrain from licking the reward ports. Licking of the reward ports resulted in resetting the 1-s interval. After the holding period, a 0.5 s stimulus was delivered as vibration to the hindlimbs applied through a flat plexiglass plate to the hindlimbs, or through electrodes implanted in the DCN or S1. A white noise auditory signal was applied over the stimulus period to mask the sound of vibration stimuli and act as a stimulus cue. A 2-s response window followed the stimulus presentation in which mice were trained to initiate licking of the left water port in response to lower frequencies and to the right water port in response to higher frequencies. To minimize a direction bias, the trial type was chosen pseudo-randomly by allowing a maximum of three trials of the same type in a row. Hit trials (correctly licking left for lower frequencies and right for higher frequencies) were rewarded by a drop of water and the immediate transition to a new trial. Miss trials (licks to the incorrect side) resulted in a timeout punishment of 2-4 seconds. If mice did not lick within the 2-s response window this resulted in a 2-s timeout. After the timeout period a new trial started with the hold period in which mice had to refrain from licking for 1-s.

Five mice (n = 5) were trained for 14-26 days on a 2AFC vibration task in which mice had to discriminate 200 Hz and 800 Hz. Once testing began, vibration stimuli were restricted to 200 Hz compared to 600 Hz for 80% of the trials and 20% of the trials were pseudo randomly switched to probe trials in which electrical stimuli of 200 Hz or 600 Hz were applied to the DCN or S1. These same five mice also completed an electrical only discrimination task in which they had to discriminate 200 Hz, 300 Hz, and 400 Hz from a 100 Hz standard. The task was the same as described above: i.e. left licks in response to low frequencies and right licks in response to high frequencies were rewarded. For each experiment, testing began when mice reached sensitivity performance of d’ > 1 over 100 trials.

Electrical stimuli consisted of 0.2 ms square pulses of 100 Hz, 200 Hz, 300 Hz, 400 Hz, or 600 Hz and amplitudes of 100-300 μA. Vibration stimuli were 200 Hz, 600 Hz or 800 Hz sinusoidal stimuli with an amplitude of 3 μm.

Mice performance was calculated as D-prime, d’ = z(hit rate) - z(miss rate), where z(p) is the inverse of the cumulative Gaussian distribution function. The behavioral task was programmed and controlled with the Bpod real-time system (Sanworks). Behavioral data were analyzed in MATLAB (Mathworks) and R (R statistical computing).

#### Focused ion beam scanning electron microscopy (FIB-SEM)

Animals were anesthetized with sodium pentobarbital (60 mg/kg) by intraperitoneal injection. They were subsequently transcardially perfused with 15 ml of 0.1M sodium phosphate buffer pH 7.4 and then 60 ml of 2% paraformaldehyde (15710-S, Electron Microscopy Sciences) 1% glutaraldehyde (G7526, Sigma-Aldrich) in 0.1M sodium phosphate buffer pH 7.4. The animals were left for two hours and then the DCN and thalamus were dissected and post-fixed overnight at 4°C in 2% paraformaldehyde 1% glutaraldehyde in 0.1M sodium phosphate buffer pH 7.4. They were then prepared for focused ion beam scanning electron microscopy.

##### Sample preparation

After overnight post-fixation the samples were prepared for FIB-SEM with support from Pilar Ruga Fahy at the PFMU Electron Microscopy Facility at the University of Geneva. The sample preparation was based on the protocol for preparation of biological specimens for volumetric electron microscopy by Ellisman (Deerinck et al. 2022; Georgiou et al. 2022). The serial sections were thoroughly rinsed (5 x 3 minutes) in 0.15M cacodylate buffer with 2mM CaCl_2_ and then postfixed in 4% osmium tetroxide with 3% potassium ferrocyanide for an hour on ice, followed by 20 minutes in 1% thiocarbohydrazide at room temperature and then 30 minutes in 2% osmium tetroxide. In between each solution the samples were thoroughly washed with ddH_2_O (5 x 3 minutes). The samples were left overnight in 1% uranyl acetate at 4°C. The next morning the samples were rinsed again (ddH_2_O, 5 x 3 minutes) and subsequently incubated in lead aspartate (0.132 g NiPb/ 20 mL aspartic acid, adjusted to pH 5.5 with 1N KOH) at 60°C. The samples were rinsed (ddH_2_O, 5 x 3 minutes) a final time before being dehydrated through increased concentrations of ethanol (30%, 50%, 70%, 90% for 7 minutes each; 2 x 100% for 30 minutes each) and then embedded in epoxy embedding medium kit (Epon substitute embedding medium kit; 45359, Sigma Aldrich). For this, the samples were left in 100% propylene oxide for 10 minutes, 25% epoxy resin 75% propylene for 30 minutes, 50% epoxy resin 50% propylene for 30 minutes, 100% epoxy resin for 30 minutes and finally 100% epoxy resin overnight. The samples were then left to harden for 48 hours in a 65°C oven.

##### Imaging

Imaging was carried out by Iryna Nikonenko from the Electron Microscopy Facility (PFMU) at the University of Geneva (Switzerland) on a Helios 660 Nanolab Dual Beam (FEI) using the Auto Slice and View (ASV 4.2) software. The resin block was trimmed down to the area of interest with a glass knife and mounted on stub pins. Semi-thin (200 nm) sections were cut on the ultramicrotome and stained with toluidine blue to confirm the region of interest. The samples were coated in 20 nm of gold. For the imaging, an acceleration voltage of 2 kV was used with a current of 0.4 nA. The ion beam was set to an acceleration voltage of 30 kV with a current of 0.79 nA. The DCN sample was imaged at a pixel size of 4 nm and slice thickness of 10 nm. The field of view was 25 by 18 µm and a total stack of 1684 slices was acquired. The thalamus sample was imaged at a pixel size of 6 nm and slice thickness of 10 nm. The field of view was 36 by 28 µm and a total of 600 slices were acquired.

##### Image alignment and segmentation

The TrakEM2 extension of Fiji (www.fiji.sc) was used to align the images acquired. The stack was then imported into the software Amira (Thermo Fisher Scientific), where the reconstructions were carried out. The different structures - synapses and vesicles - were manually segmented and labeled in the Segmentation workspace. Only synapses with large pre-synaptic elements (at least 2 µm, multiple mitochondria, often multiple synaptic contacts) that have a higher likelihood of being driver synapses were included in the study. The final reconstructions were rendered in the Project workspace using the “interactive thresholding” and “generate surface” tools. Surface area and volume were calculated from the generated surfaces using the “Surface area Volume” tool.

## Acknowledgements

We thank A. Taylor and J. Marozeau for advice and comments on the manuscript; G. Cuenu for help with histological techniques; R. Vickery for advice on electrophysiology experiments; P. Ruga Fahy for assistance with FIB-SEM sample preparation and I. Nikonenko for FIB-SEM imaging; and G. Cuenu for help with breeding mice. This work was supported by the Swiss National Science Foundation (310030_184829), the European Research Council (OPTOMOT), and the International Foundation for Research in Paraplegia. K.-S.L. is a EMBO Postdoctoral Fellow (ALTF_816-2020).

## Author contributions

K.-S.L, M.P. and D.H. conceptualized the study and designed experiments. K.-S.L. ran experiments and analyzed data, with assistance from M.S. D.T.W. collected and analyzed data on the electron microscopy. A.L. collected and analyzed data on mice behavior. K.-S.L. and D.H. wrote the manuscript with assistance from A.L. and D.T.W.

